# Novel *mutations within* PRSS1 Gene that could potentially cause hereditary pancreatitis: Using Comprehensive in silico Approach

**DOI:** 10.1101/519926

**Authors:** Mujahed I. Mustafa, Abdelrahman H. Abdelmoneim, Nafisa M. Elfadol, Soada A. osman, Tebyan A. Abdelhameed, Mohamed A. Hassan

## Abstract

**Background:** Hereditary pancreatitis (HP) is an autosomal dominant disorder with incomplete penetrance characterized by recurring episodes of severe abdominal pain often presenting in childhood. The comprehensive in silico analysis of coding SNPs, and their functional impacts on protein level, still remains unknown. In this study, *we aimed to identify the pathogenic SNPs in PRSS1* gene *by computational analysis approach.*

**Materials and Methods:** *We carried out in silico analysis of structural effect of each SNP using different bioinformatics tools to predict* Single-nucleotide polymorphisms *influence on protein structure and function*.

**Result:** *Two novel mutations out of 339 nsSNPs that are found be deleterious effect on the PRSS1 structure and function*.

**Conclusion:** *This is the first in silico analysis in PRSS1 gene, which* will be a valuable resource for future targeted mechanistic and population-based studies.

## 1. Introduction

Hereditary pancreatitis (HP) is an autosomal dominant disorder with incomplete penetrance characterized by recurring episodes of severe abdominal pain often presenting in childhood (1-5) First symptoms begin since childhood, mainly before 10 years old. Main symptoms are pancreatic pain and acute pancreatitis (>70%). CP morphological changes as pancreatic calcifications are diagnosed at a median age of 22-25 years. Exocrine and endocrine pancreatic insufficiency occurred in 34% and 26% at a median age of 29 and 38 years.(6, 7) The first family of hereditary pancreatitis was described in 1952.(8, 9) About 100 families have been documented worldwide.(4) which has been screened in different countries (3, 10-16)

HP is a progressive inflammatory disease in which pancreatic secretory parenchyma is destroyed and replaced by fibrous tissue(17), eventually leading to malnutrition and diabetes.(18) several studies shows that, Patients with HP have a markedly increased risk of pancreatic cancer compared with the general population up to 40-50%. (1, 16, 19-22) Also cigarette Smoking increases the risk factor for developing pancreatic cancer in Patients with Hereditary Pancreatitis.(23, 24)

Currently, there is no treatment for HP but it can be managed by pancreatic enzyme replacement therapy and analgesics are offered to control pain. In addition, endoscopic retrograde cholangiopancreatography (ERCP) and surgical intersections are reserved for all relevant complications. Patients should avoid tobacco use and alcohol exposure. Unfortunately, surgical removal of affected pancreatic tissue does not necessarily guarantee the patient’s long healthy life.(25-31)

The term hereditary pancreatitis has primarily been associated with mutations in the serine protease 1 gene *(PRSS1)* which has been mapped to chromosome 7 q35.(5, 20) PRSS1 is the most reported gene of hereditary pancreatitis.(1, 5-7, 15, 26, 30, 32-36) which encoding human cationic trypsinogen has been conclusively associated with autosomal dominant hereditary pancreatitis and sporadic nonalcoholic chronic pancreatitis. Most high-penetrance *PRSS1* variants increase intrapancreatic trypsin activity by stimulating trypsinogen autoactivation and/or by inhibiting chymotrypsin C-dependent trypsinogen degradation. (37) Interesting, mutations in the gene may protect one against the disease.(38, 39) In several studies, (R122H) mutation is strongly associated with hereditary pancreatitis.(6, 25, 32, 40, 41)

The main objective of this study, to identify functional SNPs within dbSNP located in coding nonsynonymous, 3-untranslated and 5-untranslated regions of *PRSS1* gene. Also to identify the most deleterious SNPs that could be used as diagnostic markers, using bioinformatics approach.(42-44)

## 2. Material and Methods

### 2.1. Data mining

The data on human *PRSS1* gene was collected from National Center for Biological Information (NCBI) web site. (45) The SNP information (protein accession number and SNP ID) of the *PRSS1* gene was retrieved from the NCBI dbSNP (http://www.ncbi.nlm.nih.gov/snp/) and the protein sequence was collected from Swiss Prot databases.(46) (http://expasy.org/)

### 2.2. Non-Synonymous SNPs Analysis

#### 2.2.1. Assessment of the Functional Impacts of Deleterious nsSNPs

##### SIFT (Sorting intolerant from tolerant)

Predicts whether an amino acid substitution affects protein function based on the degree of conservation of amino acid residues in sequence alignments derived from closely related sequences.(47) SIFT scores <0.05 are predicted by the algorithm to be intolerant or deleterious amino acid substitutions, whereas scores >0.05 are considered tolerant. It is available at (http://sift.bii.a-star.edu.sg/).

##### Polyphen-2

It is an online tool (48) to predict possible impact of an amino acid substitution on both structure and function of a human protein by analysis of multiple sequence alignment and protein 3D structure, in addition it calculates position-specific independent count scores (PSIC) for each of two variants, and then calculates the PSIC scores difference between two variants. The higher a PSIC score difference, the higher the functional impact a particular amino acid substitution is likely to have. Prediction outcomes could be classified as probably damaging, possibly damaging or benign according to the value of PSIC as it ranges from (0_1); values closer to zero considered benign while values closer to 1 considered probably damaging and also it can be indicated by a vertical black marker inside a color gradient bar, where green is benign and red is damaging. nsSNPs that predicted to be intolerant by Sift has been submitted to Polyphen as protein sequence in FASTA format that obtained from UniproktB /Expasy after submitting the relevant ensemble protein (ESNP) there, and then we entered position of mutation, native amino acid and the new substituent for both structural and functional predictions. PolyPhen version 2.2.2 is available at (http://genetics.bwh.harvard.edu/pph2/index.shtml).

##### Provean

Provean is an online tool (49) that predicts whether an amino acid substitution has an impact on the biological function of a protein grounded on the alignment-based score. The score measures the change in sequence similarity of a query sequence to a protein sequence homolog between without and with an amino acid variation of the query sequence. If the PROVEAN score <−2.5, the protein variant is predicted to have a “deleterious” effect, while if the PROVEAN score is >-2.5, the variant is predicted to have a “neutral” effect. It is available at (https://rostlab.org/services/snap2web/).

##### SNAP2

Functional effects of mutations are predicted with SNAP2 (50) is a trained classifier that is based on a machine learning device called “neural network”. It distinguishes between effect and neutral variants/non-synonymous SNPs by taking a variety of sequence and variant features into account. The most important input signal for the prediction is the evolutionary information taken from an automatically generated multiple sequence alignment. Also structural features such as predicted secondary structure and solvent accessibility are considered. If available also annotation (i.e. known functional residues, pattern, regions) of the sequence or close homologs are pulled in. In a cross-validation over 100,000 experimentally annotated variants, SNAP2 reached sustained two-state accuracy (effect/neutral) of 82% (at an AUC of 0.9). In our hands this constitutes an important and significant improvement over other methods. It is available at (https://rostlab.org/services/snap2web/).

#### 2.2.2. Prediction of Disease Related Mutations SNPs

##### SNPs&GO

SNPs&GO (51) is a support vector machine (SVM) based on the method to accurately predict the mutation related to disease from protein sequence. The input is the FASTA sequence of the whole protein, the output is based on the difference among the neutral and disease related variations of the protein sequence. The RI (reliability index) with value of greater than 5 depicts the disease related effect caused by mutation on the function of parent protein. The PHD SNP and PANTHER, algorithms were also used in the display of output.

##### P-Mut

PMUT a web-based tool (52) for the annotation of pathological variants on proteins, allows the fast and accurate prediction (approximately 80% success rate in humans) of the pathological character of single point amino acidic mutations based on the use of neural networks. It is available at (http://mmb.irbbarcelona.org/PMut).

## 3. Protein Stability Analysis

### I-Mutant 3.0

I-Mutant 3.0 Is a neural network based tool for the routine analysis of protein stability and alterations by taking into account the single-site mutations. The FASTA sequence of protein retrieved from UniProt is used as an input to predict the mutational effect on protein stability.(53) It is available at (http://gpcr2.biocomp.unibo.it/cgi/predictors/I-Mutant3.0/I-Mutant3.0.cgi).

### MUpro

MUpro is a support vector machine-based tool for the prediction of protein stability changes upon nonsynonymous SNPs. The value of the energy change is predicted, and a confidence score between −1 and 1 for measuring the confidence of the prediction is calculated. A score <0 means the variant decreases the protein stability; conversely, a score >0 means the variant increases the protein stability.(54) It is available at (

### Identification of Functional SNPs in Conserved Regions by using ConSurf server

ConSurf web server provides evolutionary conservation profiles for proteins of known structure in the PDB. Amino acid sequences similar to each sequence in the PDB were collected and multiply aligned using CSI-BLAST and MAFFT, respectively. The evolutionary conservation of each amino acid position in the alignment was calculated using the Rate 4Site algorithm, implemented in the ConSurf web server. The algorithm takes explicitly into account the phylogenetic relations between the aligned proteins and the stochastic nature of the evolutionary process. Rate 4 Site assigns a conservation level for each residue using an empirical Bayesian inference. Visual inspection of the conservation patterns on the 3-dimensional structure often enables the identification of key residues that comprise the functionally-important regions of the protein.(55, 56) It is available at (http://consurf.tau.ac.il/).

### BioEdit

BioEdit is an easy-to-use biological sequence alignment editor. This free software is intended to supply a single program that can handle most simple sequence and alignment editing and manipulation functions as well as a few basic sequences analyses.(57) It is available for download at (http://www.mbio.ncsu.edu/bioedit/bioedit.html).

## 4. GeneMANIA

We submitted genes and selected from a list of data sets that they wish to query. GeneMANIA approach to know protein function prediction integrate multiple genomics and proteomics data sources to make inferences about the function of unknown proteins.(58) It is available at (http://www.genemania.org/)

## 5. Structural Analysis

### 5.1. Detection of nsSNPs Location in Protein Structure

Mutation3D is a functional prediction and visualization tool for studying the spatial arrangement of amino acid substitutions on protein models and structures. Mutation3D is able to separate functional from nonfunctional mutations by analyzing a combination of 8,869 known inherited disease mutations and 2,004 SNPs overlaid together upon the same sets of crystal structures and homology models. Further, it present a systematic analysis of whole-genome and whole-exome cancer datasets to demonstrate that mutation3D identifies many known cancer genes as well as previously underexplored target genes(59). It is available at (http://mutation3d.org).

### 5.2. Modeling nsSNP locations on protein structure

Project hope is a new online web-server to search protein 3D structures (if available) by collecting structural information from a series of sources, including calculations on the 3D coordinates of the protein, sequence annotations from the UniProt database, and predictions by DAS services. Protein sequences were submitted to project hope server in order to analyze the structural and conformational variations that have resulted from single amino acid substitution corresponding to single nucleotide substitution. It is available at (http://www.cmbi.ru.nl/hope).

### 5.3. Modeling Amino Acid Substitution

UCSF Chimera is a highly extensible program for interactive visualization and analysis of molecular structures. Chimera (version 1.8) software was used to scan the 3D (three-dimensional) structure of specific protein, and hence modifies the original amino acid with the mutated one to see the impact that can be produced. The outcome is then a graphic model depicting the mutation.(60) Chimera (version 1.8) available at (http://www.cgl.ucsf.edu/chimera/).

## 6. Results

## 7. Discussion

Two novel mutations have been found *(see table 3) that* effect on the stability and function of the *PRSS1* gene using bioinformatics tools. The methods used were based on different aspects and parameters describing the pathogenicity and provide clues on the molecular level about the effect of mutations. It was not easy to predict the pathogenic effect of SNPs using single method. Therefore, multiple methods were used to compare and rely on the results predicted. In this study we used different in silico prediction algorithms: SIFT, PolyPhen-2, Provean, SNAP2, SNP&GO, PHD-SNP, PANTHER, P-MUT and I-Mutant 3.0, MUpro, ConSurf and Mutation3D.

Single-nucleotide polymorphism (SNP) association studies have become crucial in revealing the genetic correlations of genomic variants with complex diseases. In silico analysis has been done for many disorders for cancer related genes and other disorders. (61-63) Therefore, we used deep comprehensive in silico analysis to study nsSNP in Homo sapiens located in coding region of *PRSS1* gene, were investigate in dbSNP/NCBI Database, out of 911 there are 339 nsSNPs (missense mutations),(see figure 1) were submitted to SIFT server, PolyPhen-2 server, Provean sever and SNAP2 respectively, 141 SNPs were predicted to be deleterious in SIFT server. In PolyPhen-2 server, the result showed that were found to be damaging (54 possibly damaging and 116 probably damaging showed deleterious). In Provean server our result showed that 203 SNPs were predicted to be deleterious. While in SNAP2 server the result showed that 172 SNPs were predicted to be Effect. The differences in prediction capabilities refer to the fact that every prediction algorithm uses different sets of sequences and alignments. In table (2) we were submitted four positive results from SIFT, PolyPhen-2, Provean and SNAP2 *(see table 1)* to observe the disease causing one by SNP&GO, PHD-SNP, P-Mut and MUpro servers.

**Figure 1:**
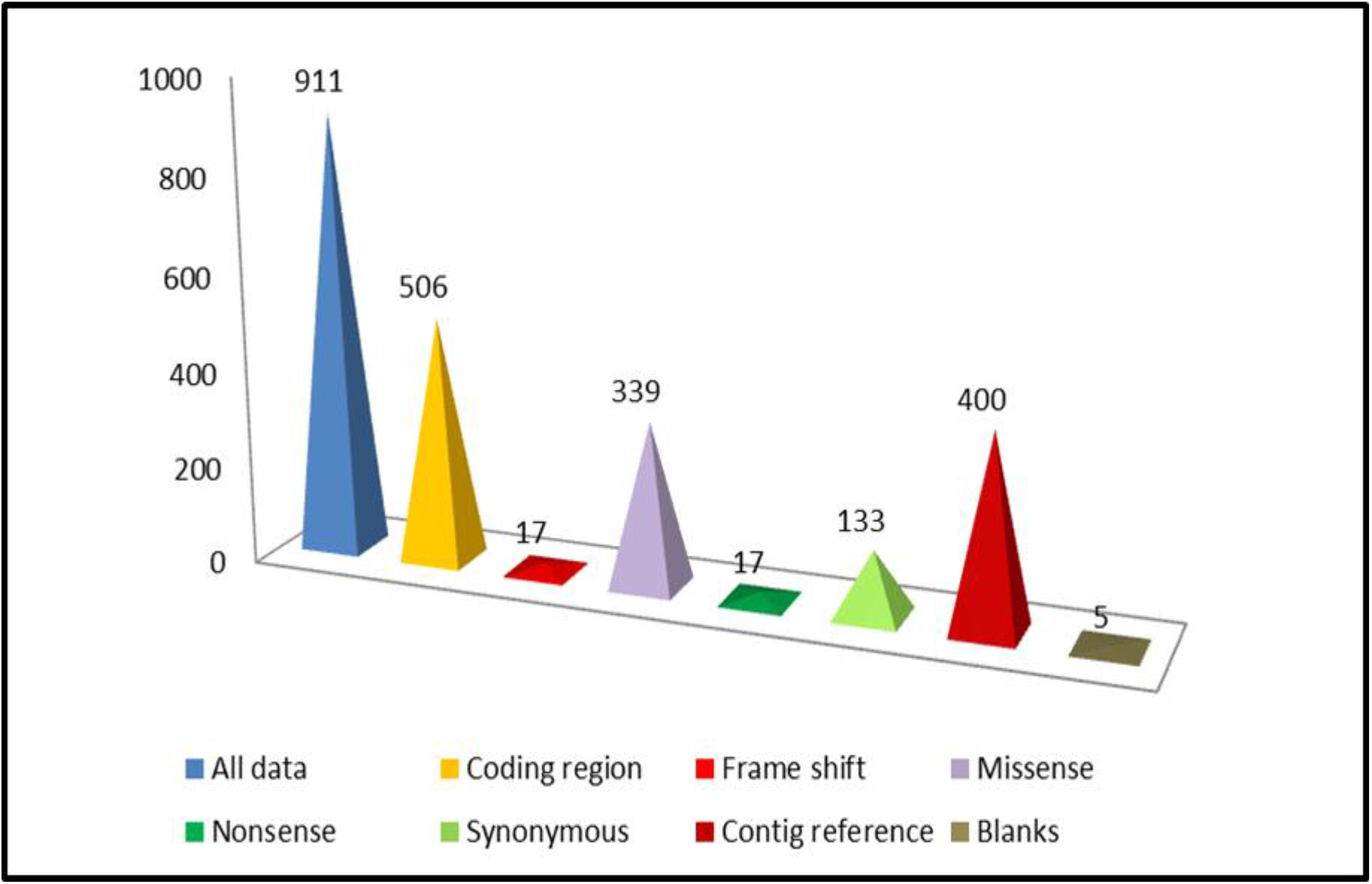
A graphical representation of the distribution of frame shift, missense, nonsense and synonymous SNPs for *PRSS1* gene in coding region (based on the dbSNP database). *Note: the numbers represent in the chart, is numbers of SNPs.

**Table (1):**
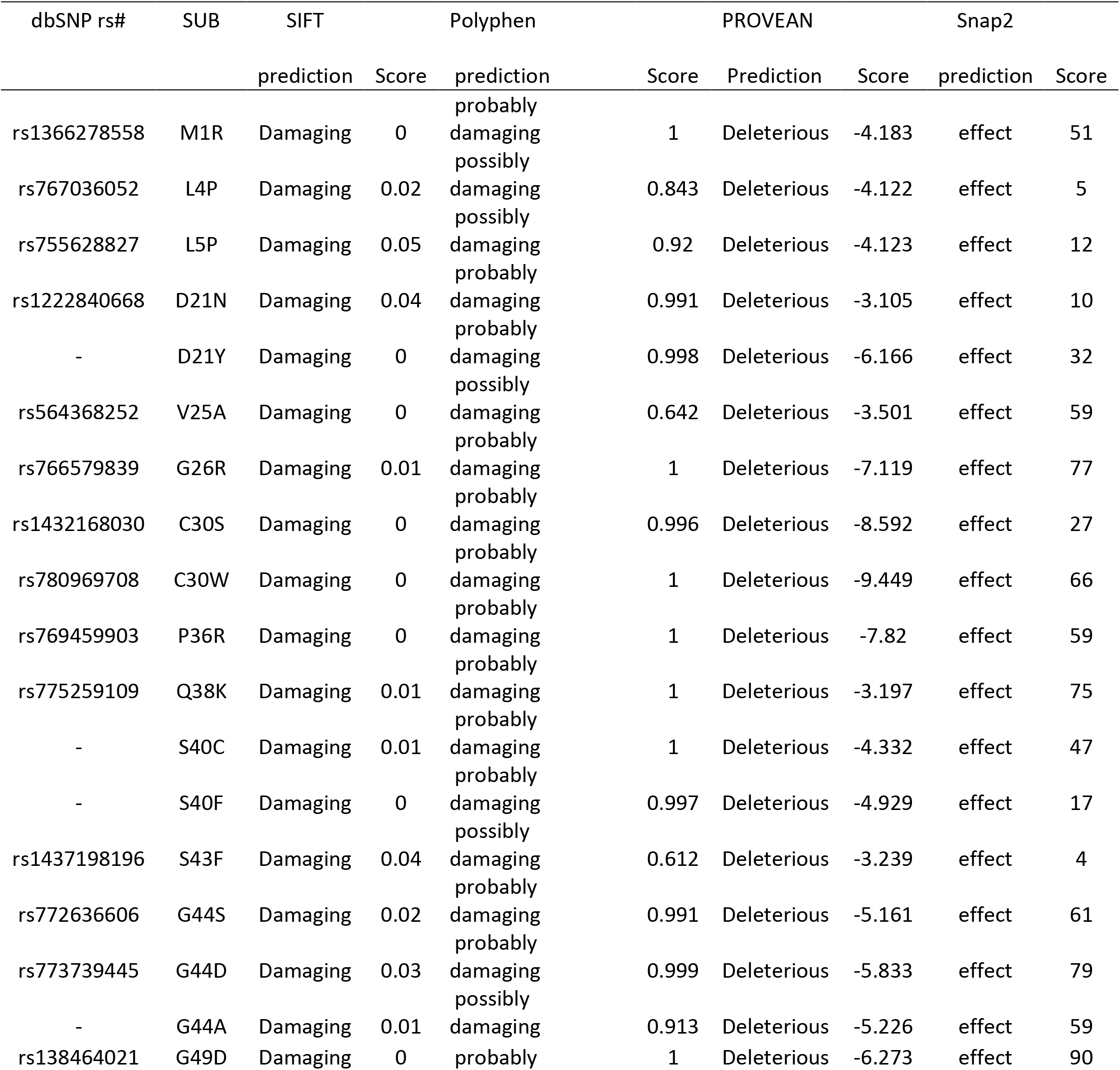
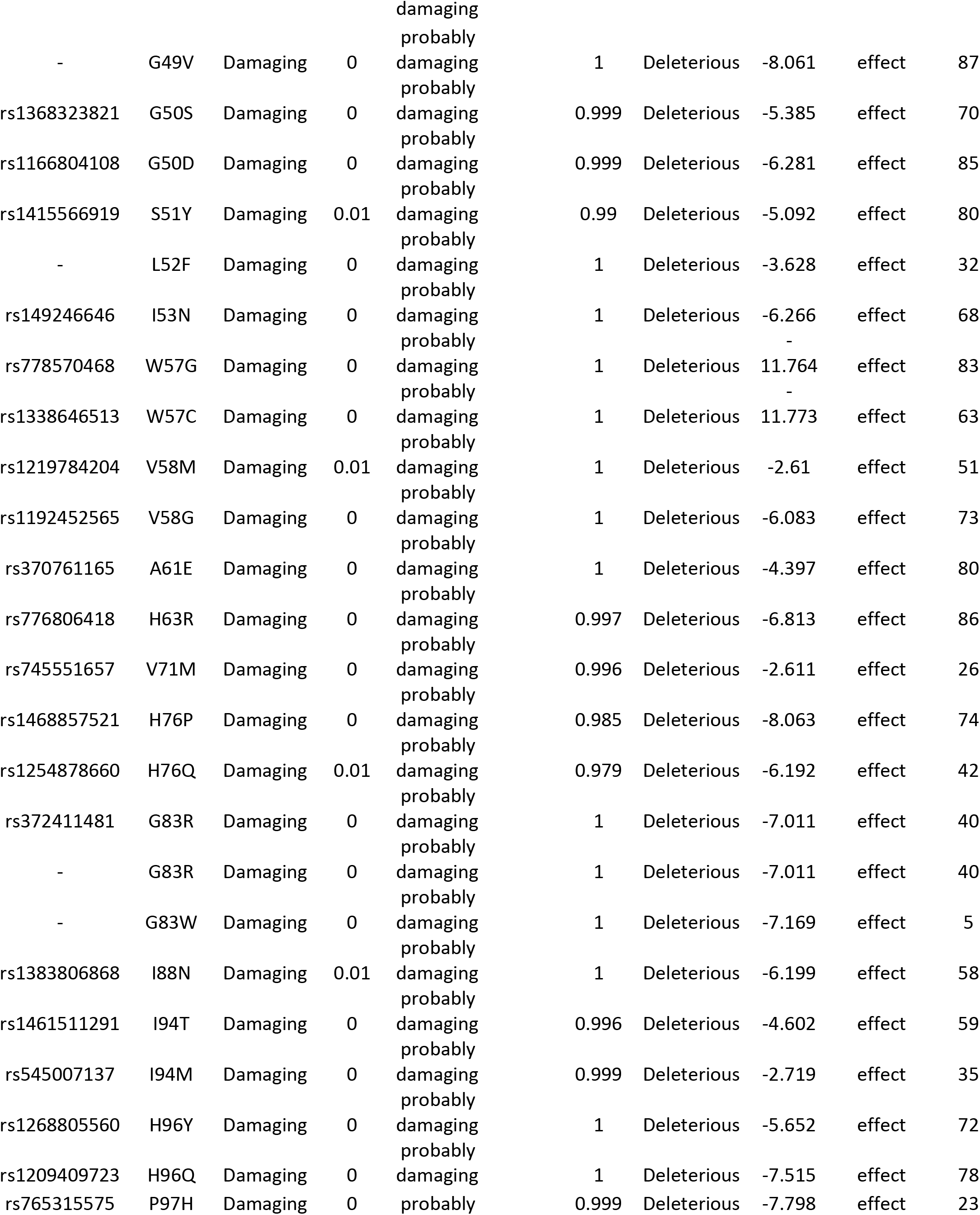
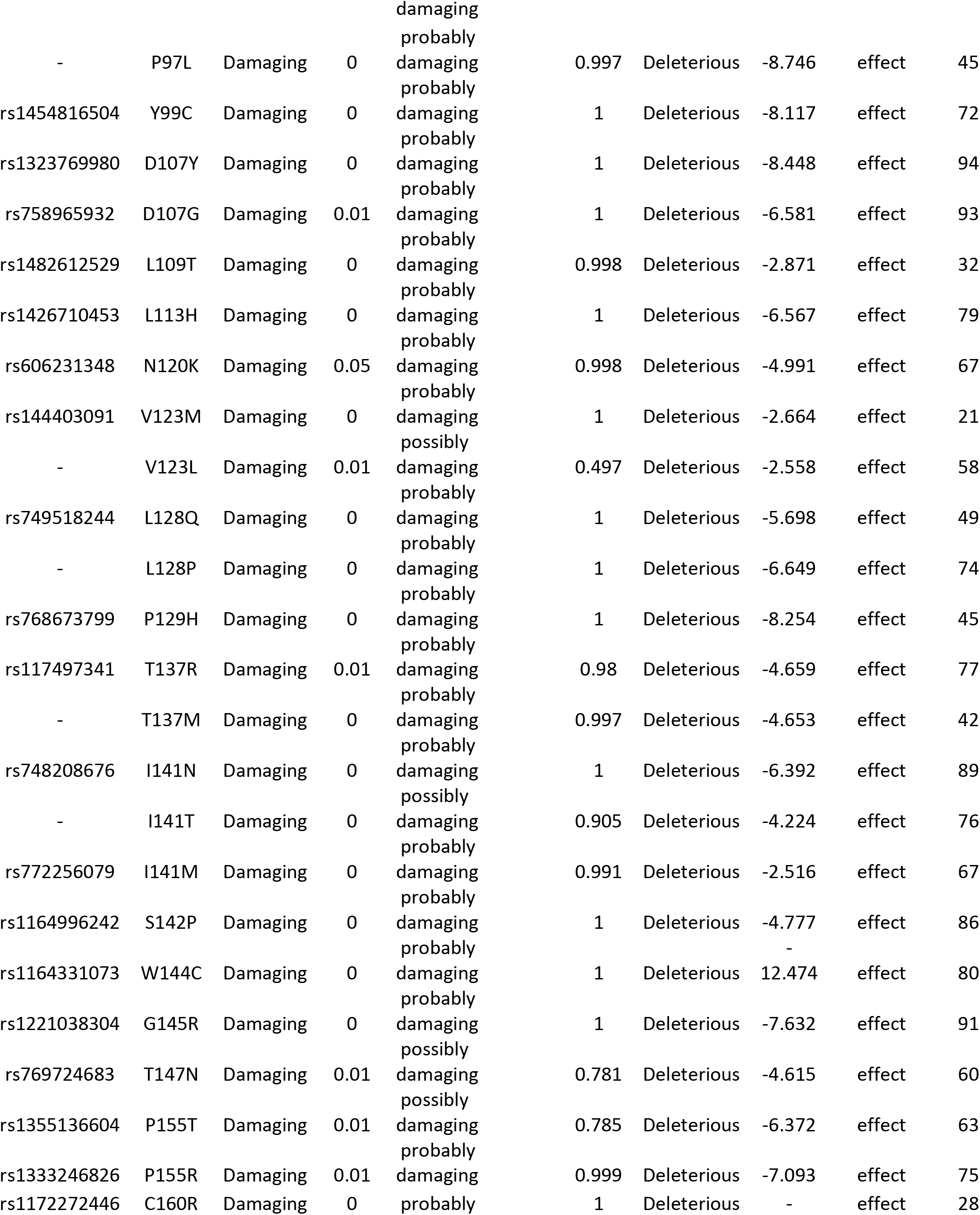
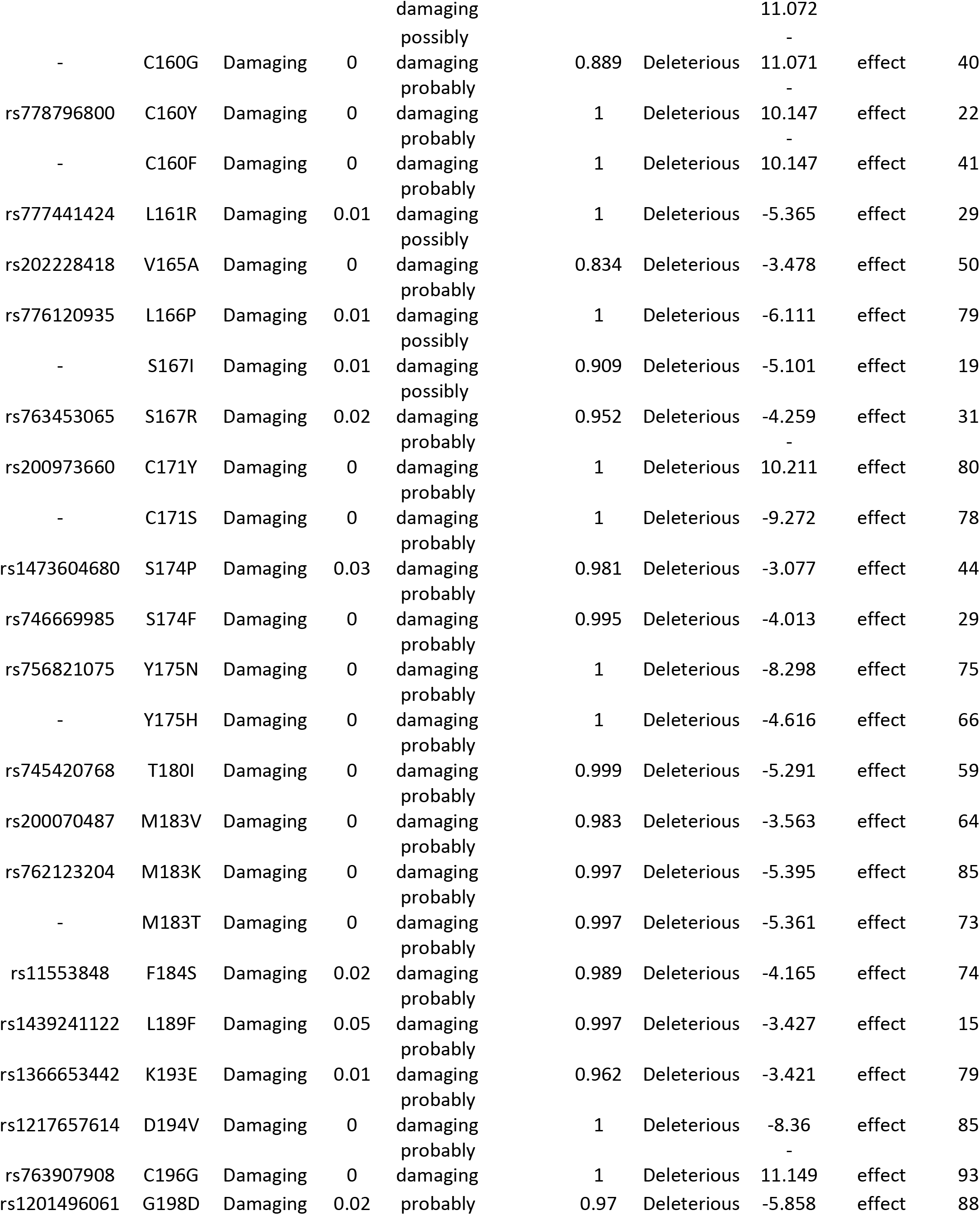
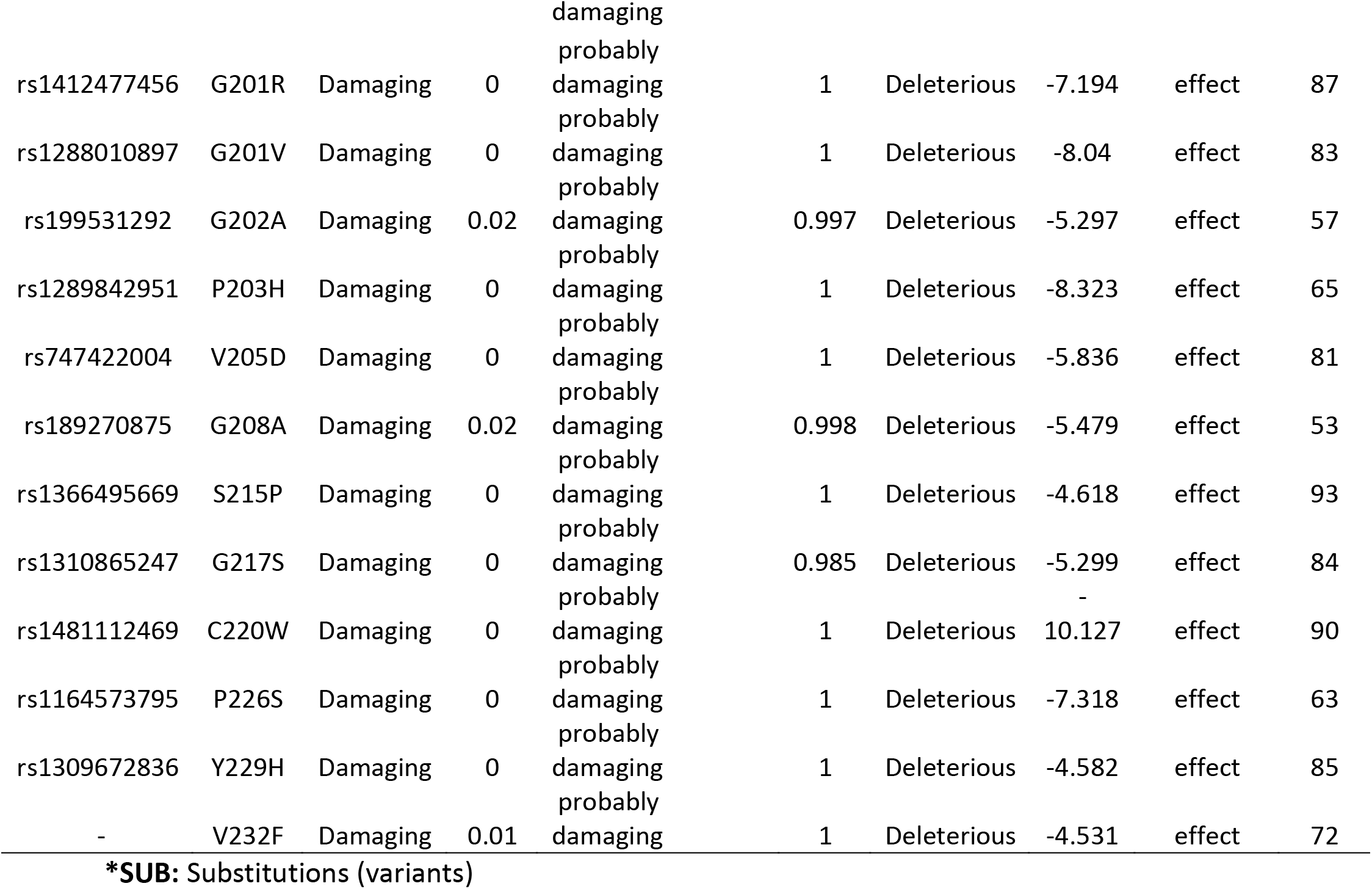
Damaging or Deleterious or effect nsSNPs associated variations predicted by various online tools.

**Table (2):**
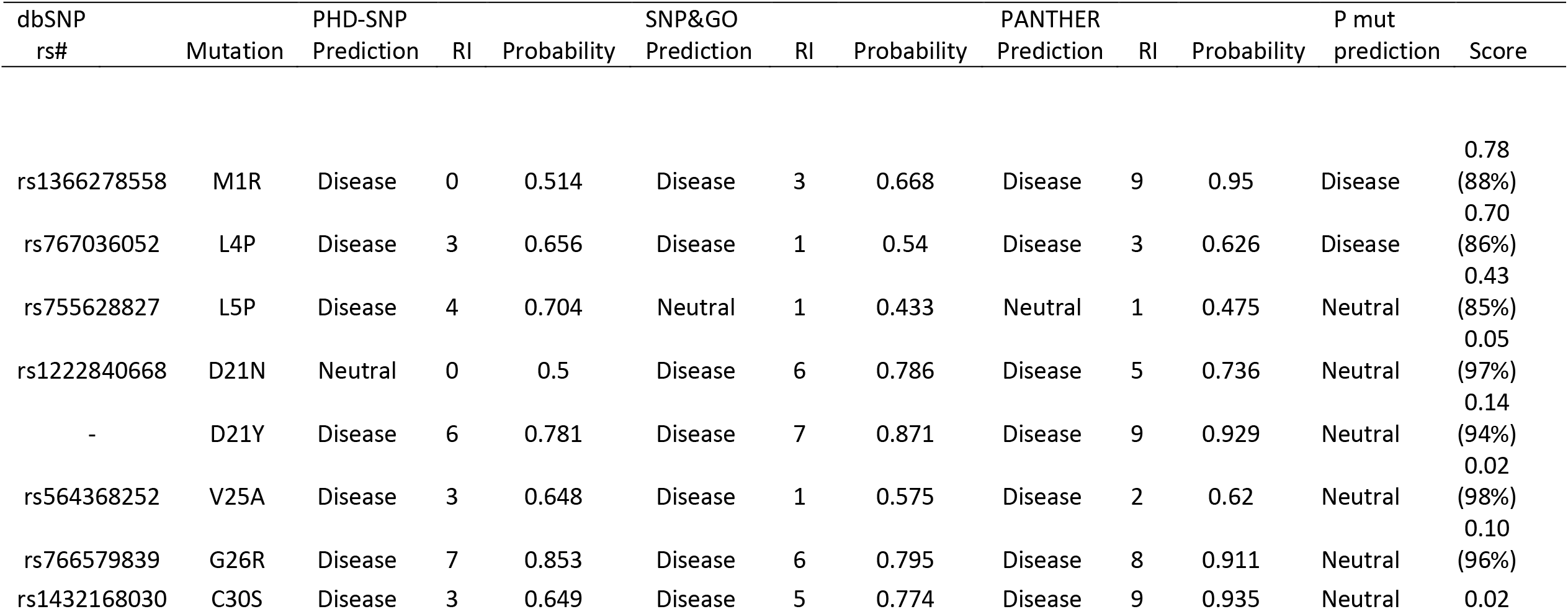
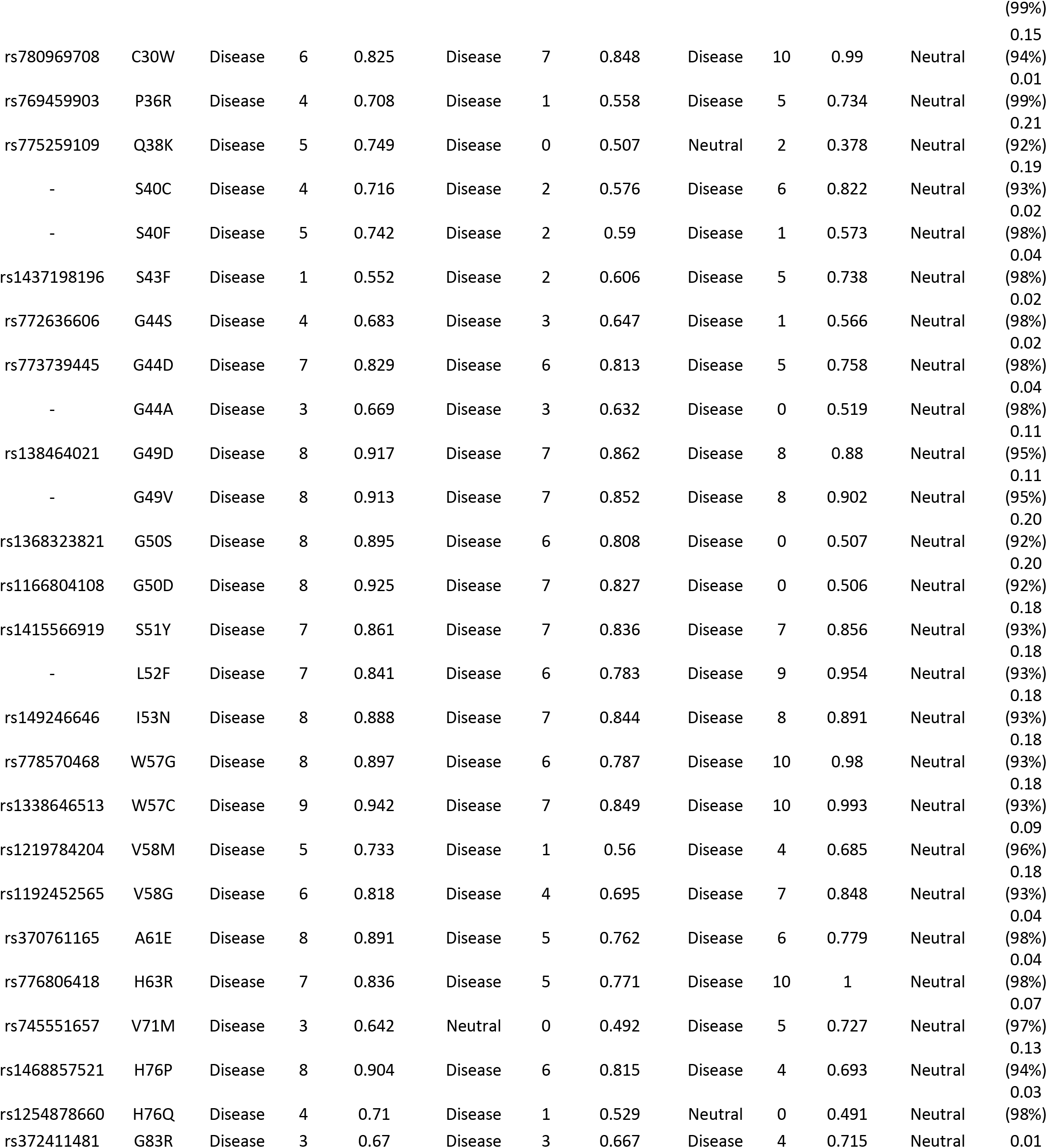
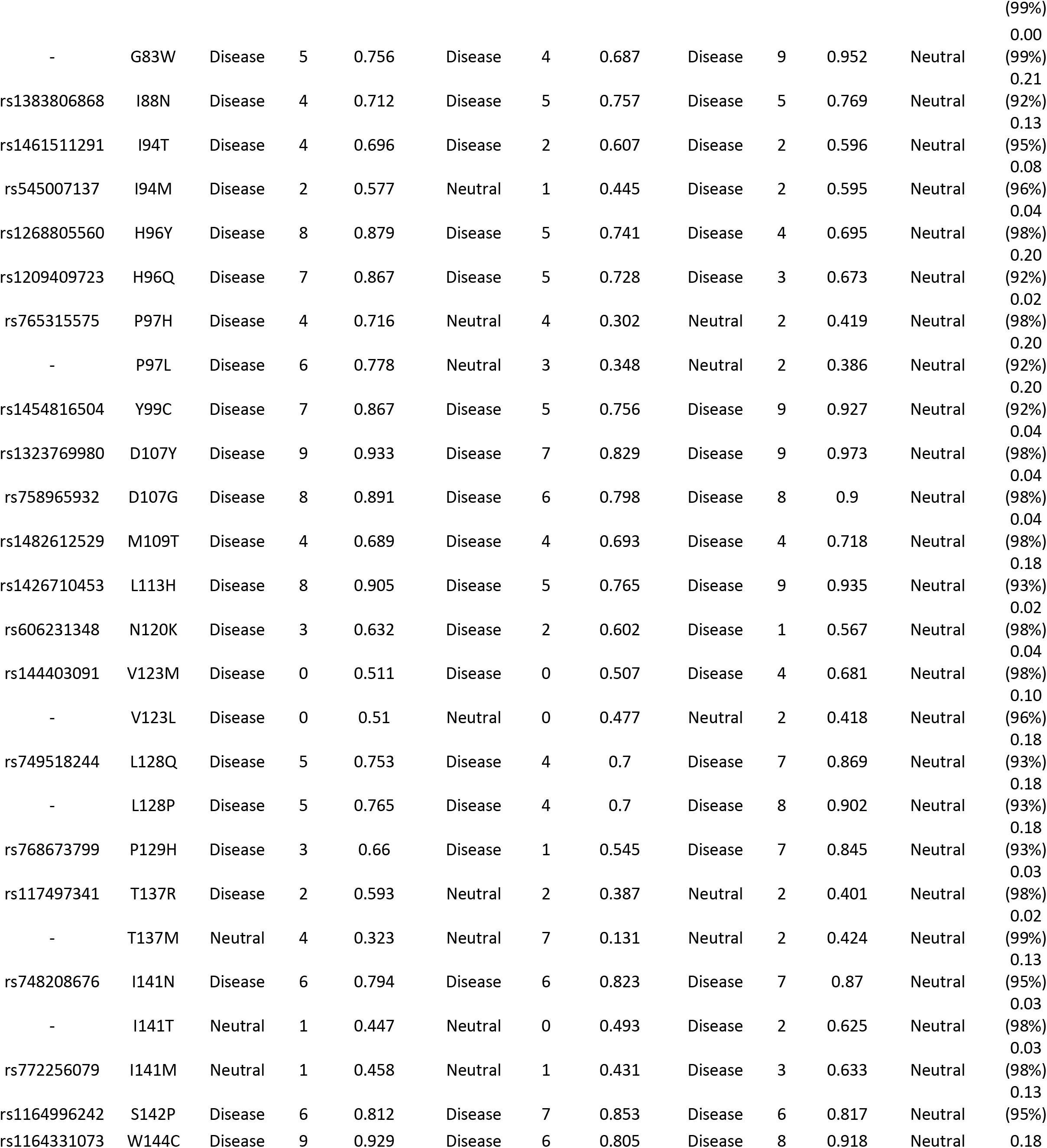
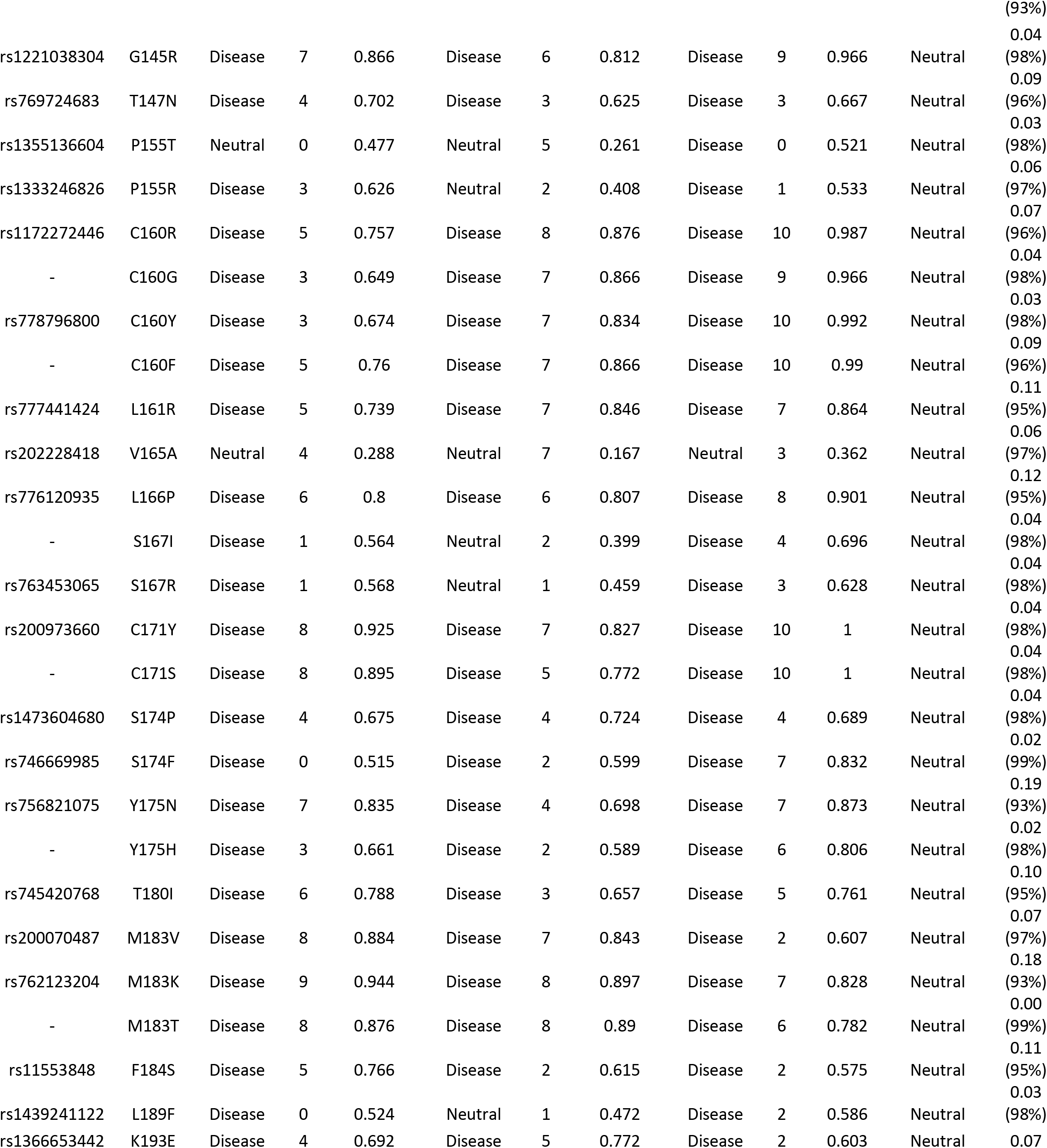
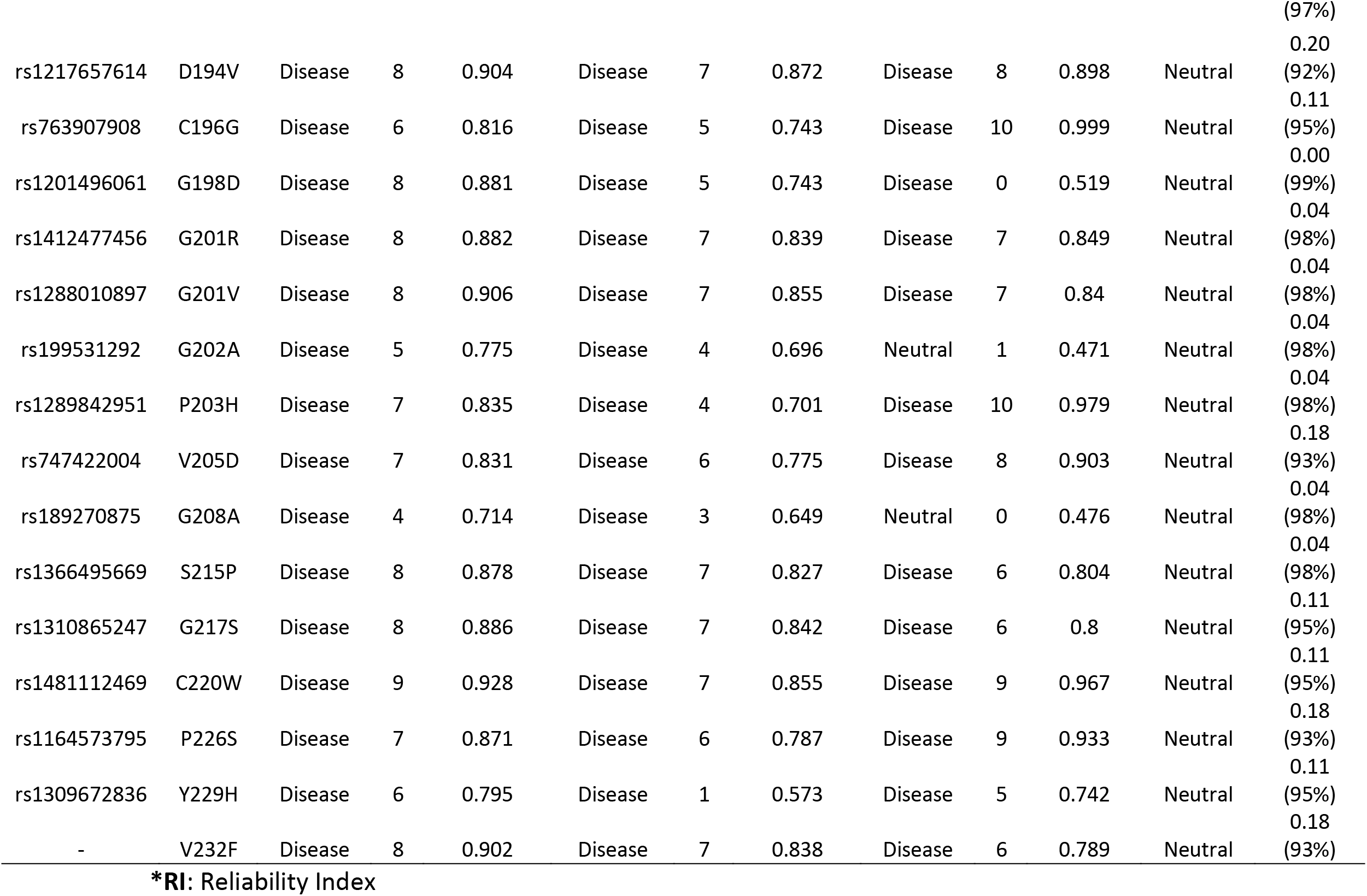
List of non-synonymous SNPs analyzed for disease association by SNP&GO, PHD-SNP, PANTHER and P-mut.

**Table (3):** Protein Stability Analysis predicted by I-Mutant v3.0 and Mupro (also Show the two novel mutations)

| dbSNP rs# | Amino Acid change | SVM2 Prediction Effect | RI | DDG Value Prediction | Mupro Prediction | Mupro Score |
| --- | --- | --- | --- | --- | --- | --- |
| rs1366278558 | M1R | Decrease | 1 | -0.62 | Decrease | -1.018261 |
| rs767036052 | L4P | Decrease | 8 | -1.66 | Decrease | -1.623945 |

\*RI: Reliability Index

In SNP&GO, PHD-SNP, PANTHER and P-Mut servers were used to predict the association of SNPs with disease. According to SNP&GO, PHD-SNP, PANTHER and P-Mut (85, 95, 90 and 2 SNPs respectively) were found to be disease related SNPs. We selected the four positive results (disease related SNPs) for further analysis by I-Mutant 3.0 and MUpro, Table (3) (figure 2) While I-Mutant and MUpro results revealed that the protein stability decreased which destabilize the amino acid interaction, (see Table 3).

**Figure 2:**
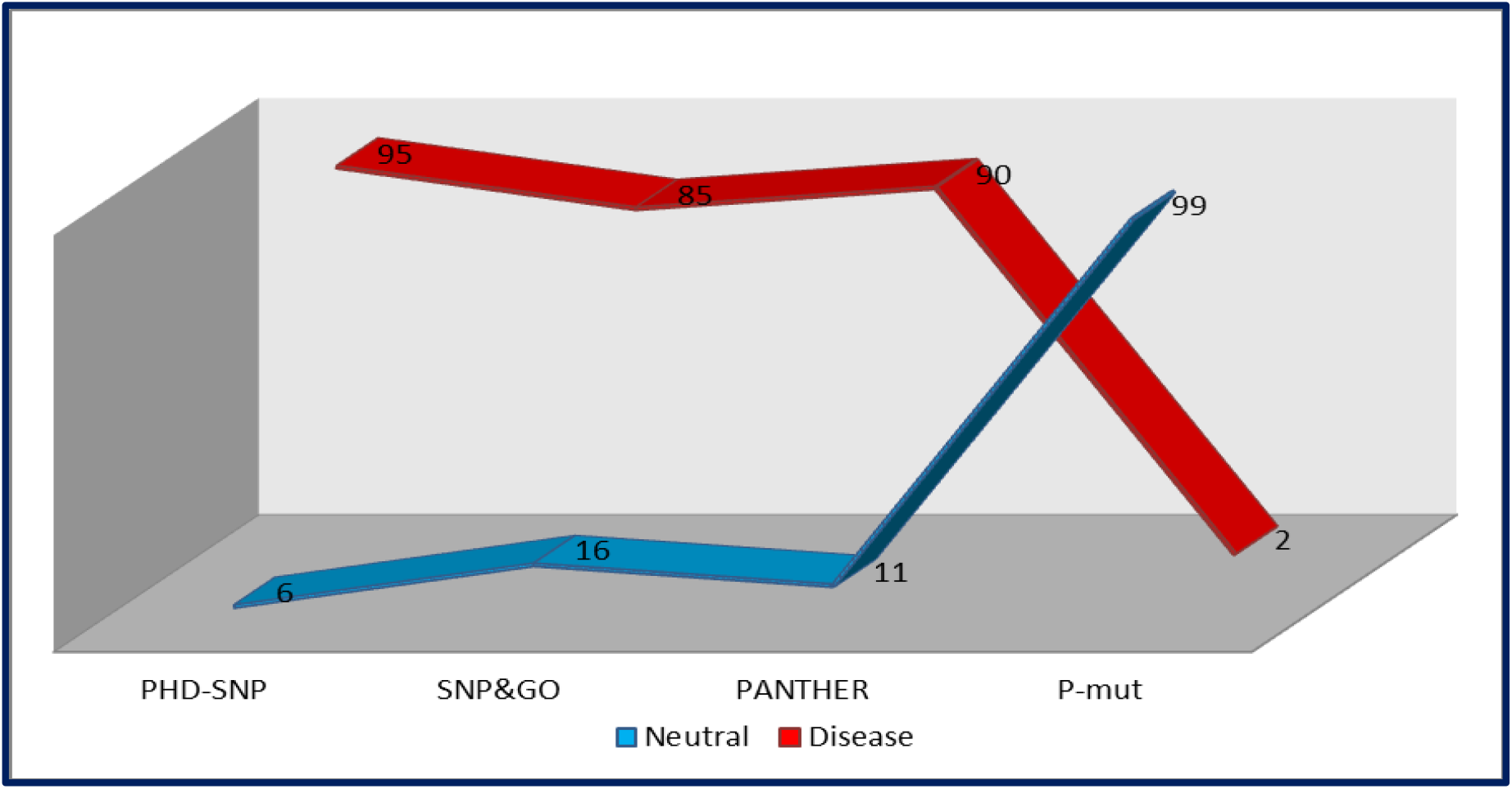
Graphical representation of Disease variations by PHD-SNP, SNP&GO, PANTHER and P-mut.

GeneMANIA revealed that *PRSS1* has many vital functions: blood microparticle, cobalamin metabolic process, extracellular matrix organization, extracellular structure organization,serine hydrolase activity, serine-type endopeptidase activity, serine-type peptidase activity. The genes co-expressed with, share similar protein domain, or participate to achieve similar function were illustrated by GeneMANIA and shown in figure 3, Tables (5&6).

**Figure 3:**
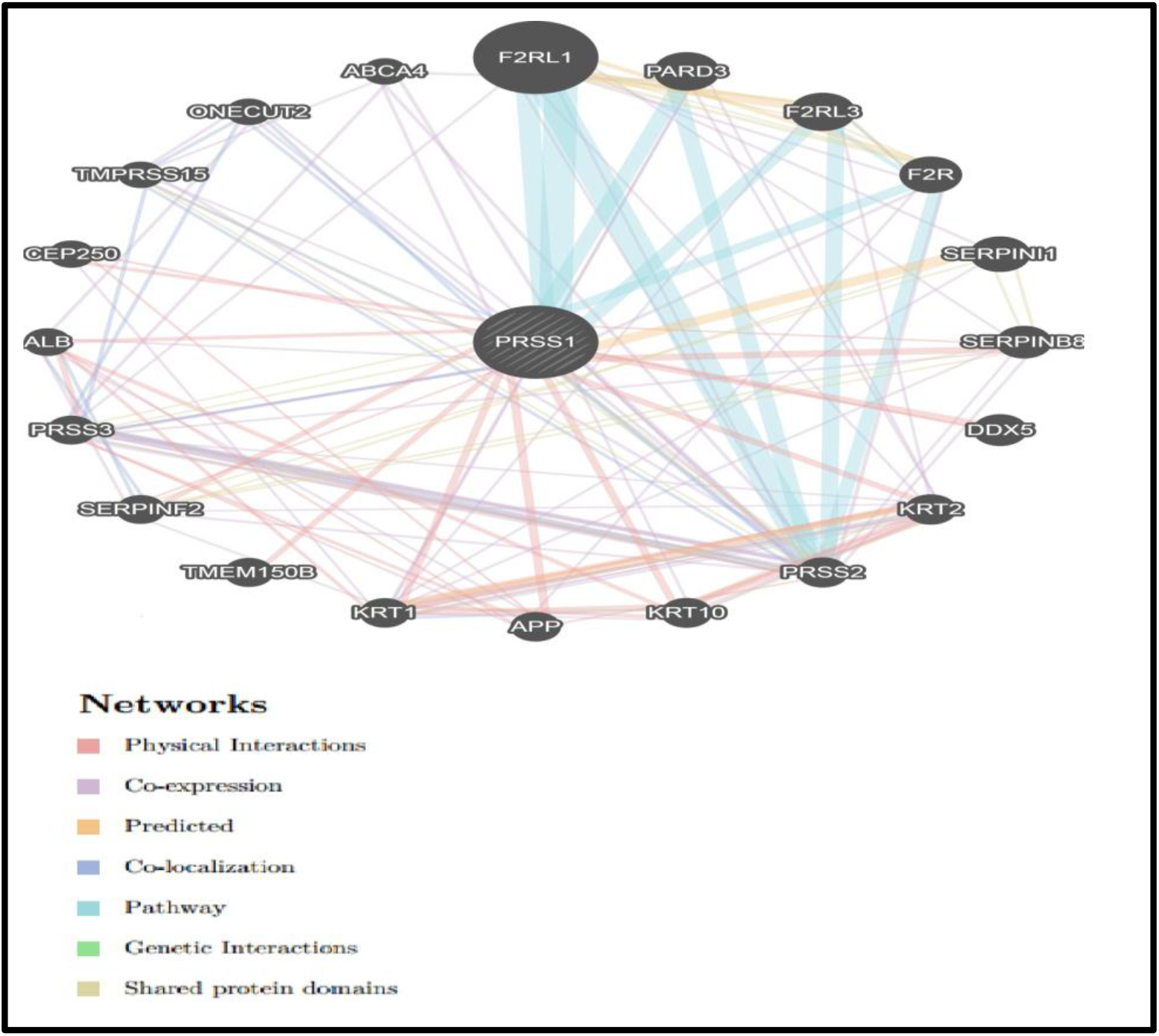
Interaction between PRSS1 and its related genes.

Project HOPE server was used to submit the two deleterious nsSNPs: (rs1366278558): (M1R): Methionine residue changed to Arginine at position 1. (Figure 4) shows the schematic structures and homology modeling of the original and mutant amino acid. The backbone, which is the same for each amino acid, is colored red. The side chain, unique for each amino acid, is colored black. Each amino acid has its own specific size, charge, and hydrophobicity value. The mutation is located within the signal peptide. This sequence of this peptide is important because it is recognized by other proteins and often cleaved of to generate the mature protein. The new residue that is introduced in the signal peptide differs in its properties from the original one. It is possible that this mutation disturbs recognition of the signal peptide. The mutant residue is smaller than the wild-type, and this will cause a possible loss of external interactions. The wild-type residue is more hydrophobic than the mutant residue and this mutation might cause loss of hydrophobic interactions with other molecules on the surface of the protein and thus may cause function loss or change.

**Figure 4:**
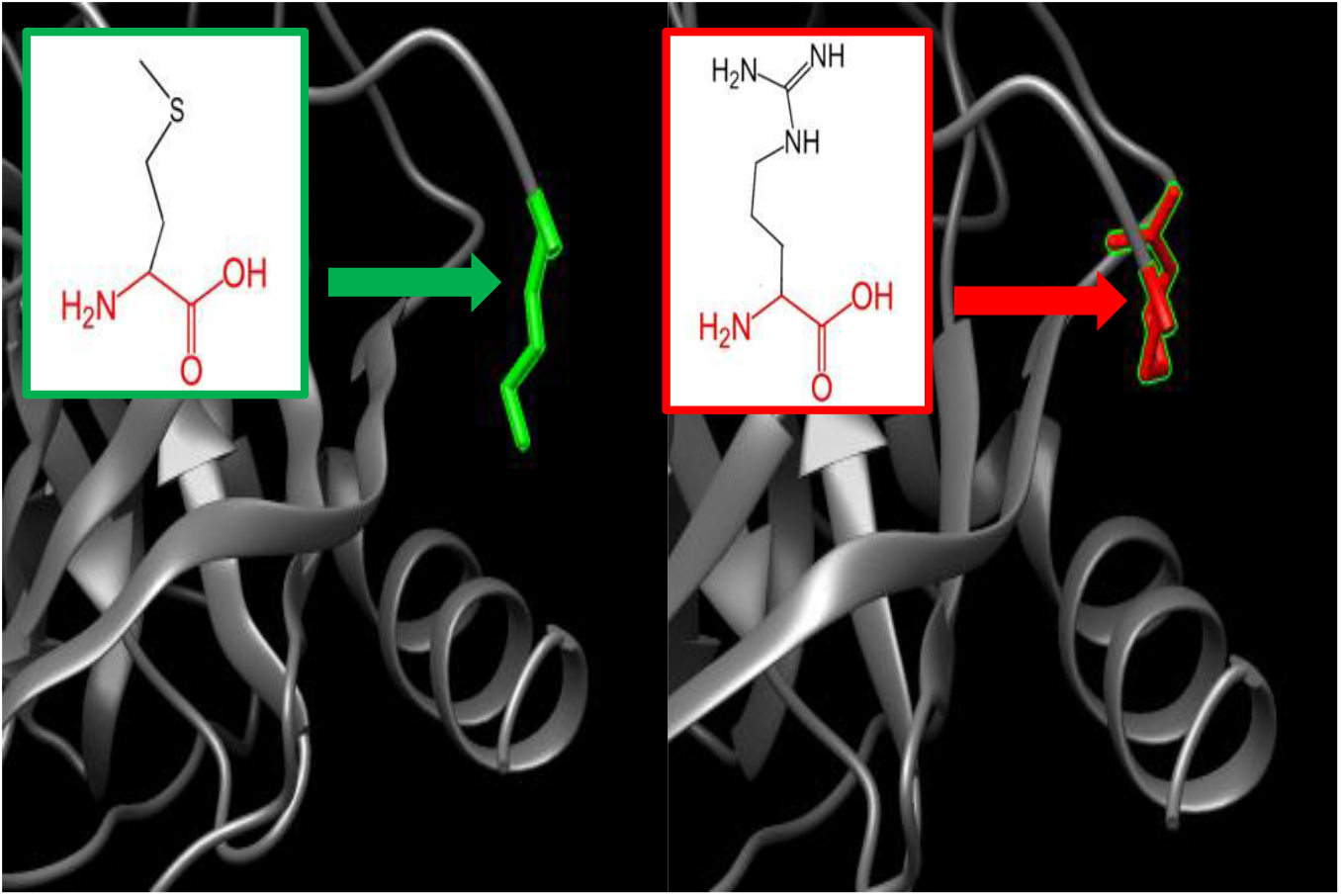
The amino acid Methionine change to Arginine at position 1.

(rs767036052):(L4P): Leucine residue changed to Proline at position 4. (Figure 5) shows the schematic structures of the original and the mutant amino acid. The backbone, which is the same for each amino acid, is colored red. The side chain, unique for each amino acid, is colored black. The mutant residue is smaller and more hydrophobic than the wild-type residue. The mutation is located within the signal peptide. This sequence of this peptide is important because it is recognized by other proteins and often cleaved of to generate the mature protein. While the new residue that is introduced in the signal peptide differs in its properties from the original one. It is possible that this mutation disturbs recognition of the signal peptide. The wild-type and mutant amino acids differ in size. The mutant residue is smaller; this might lead to loss of interactions.

**Figure 5:**
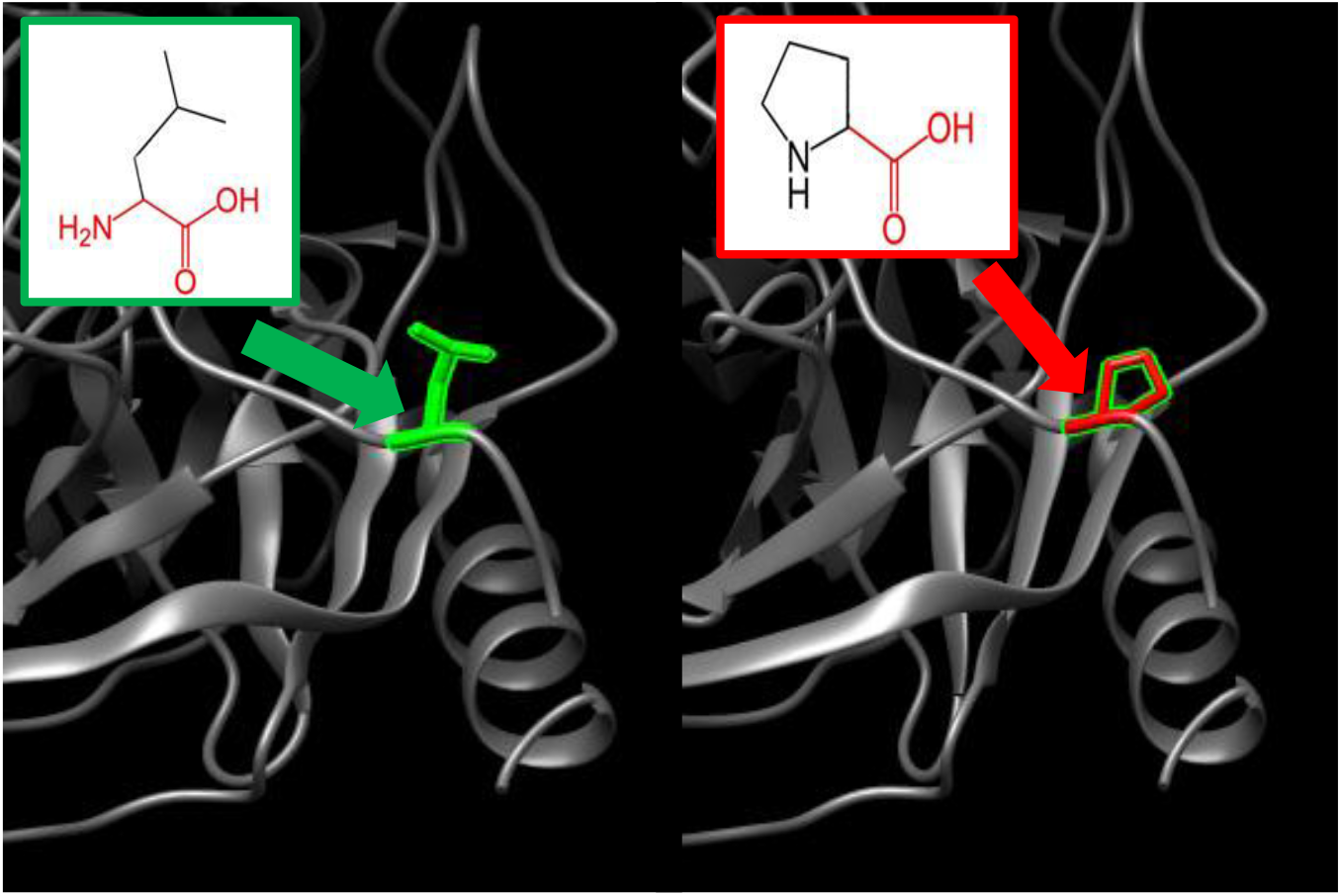
The amino acid Leucine change to Proline at position 4.

We also used ConSurf web server; the nsSNPs that are located at highly conserved amino acid positions tend to be more deleterious than nsSNPs that are located at non-conserved sites. Our ConSurf analysis revealed that (L4P) mutation was found in the highly conserved region and predicted to have a potential impact on *PRSS1* protein.(see table 4) To confirm our findings (M1R & L4P) mutations we used BioEdit (version 7.2.5) Alignment of 10 amino acid sequences of *PRSS1* confirm that, the residues predicted to be mutated are evolutionarily conserved across species. (See figure 6) Additionally, we performed analysis by Mutation3D, our result show that: (L4P) located in the domain. (See figure 7)

**Figure 6:**
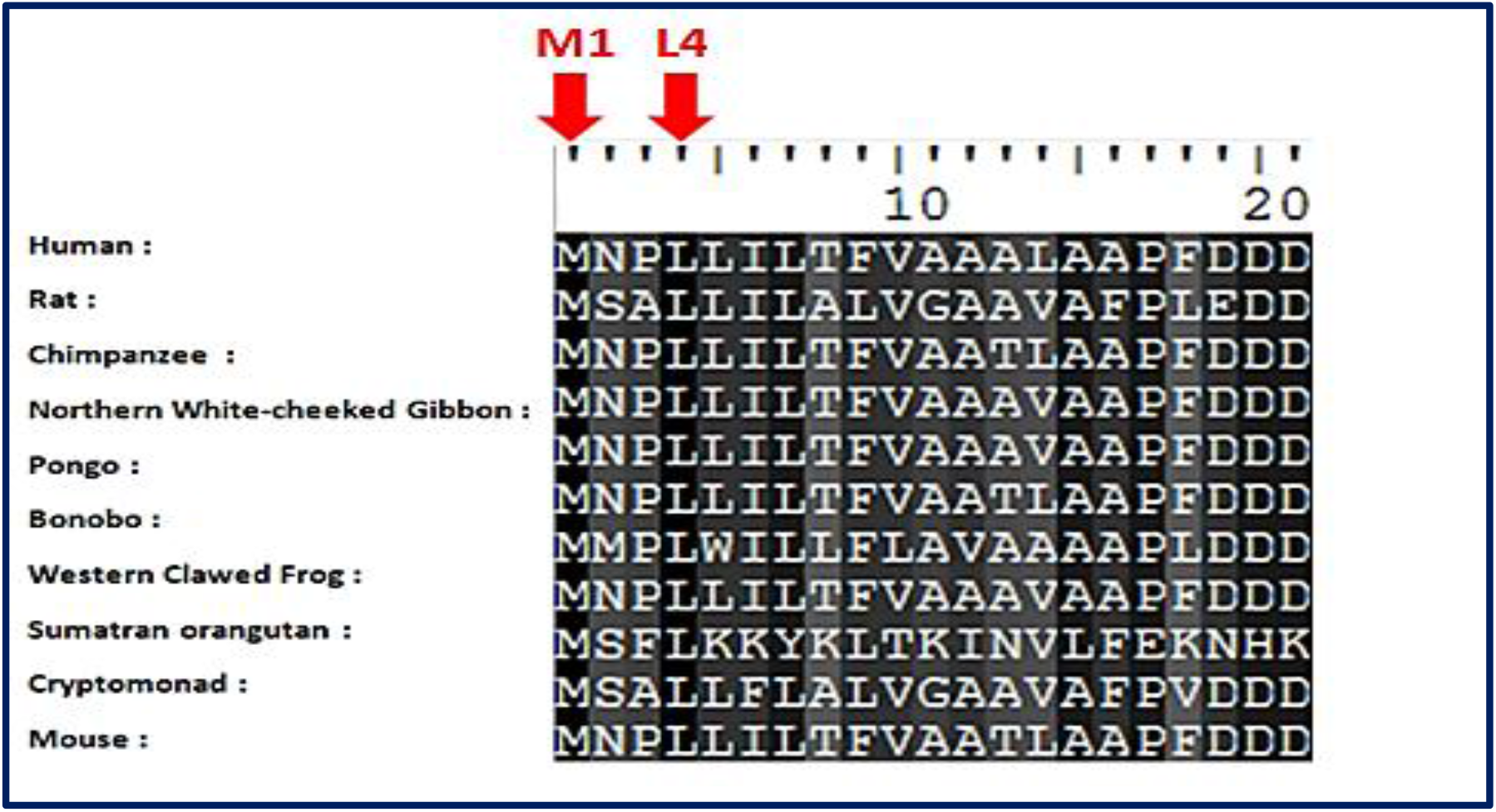
Alignment of 10 amino acid sequences of PRSS1 demonstrating that the residues predicted to be mutated (indicated by red arrow) are evolutionarily conserved across species. Sequences Alignment were done by BioEdit (version 7.2.5)

**Figure 7:**
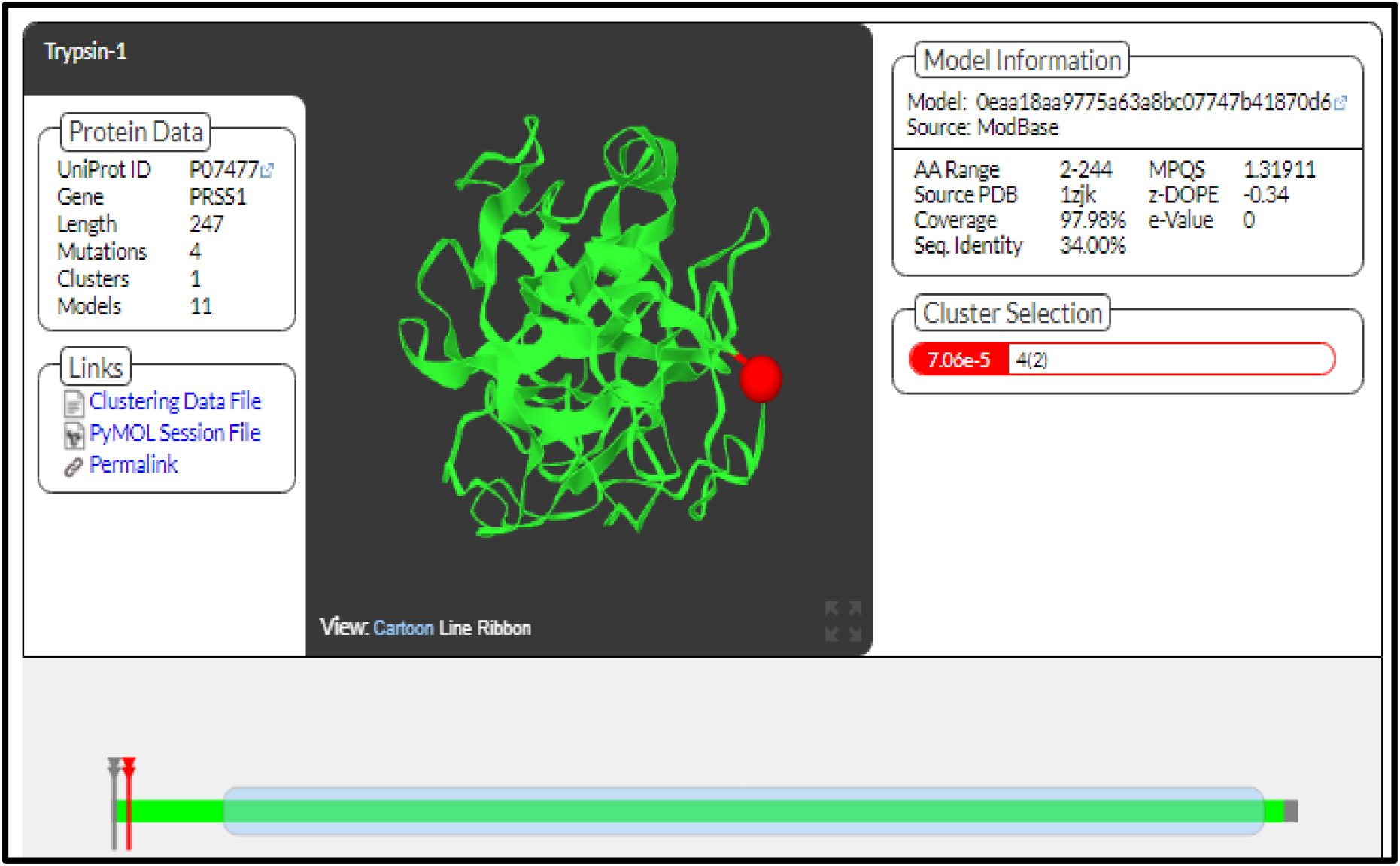
Structural models for wild type *PRSS1* and (L4P) high-risk nsSNP in its domain.

**Table (4):** Conservation profile of amino acids in *PRSS1* that coincide in location with high-risk nsSNPs.

| Residues position | Residues | CS Score normalized | Color | Residues Variety | B/E |
| --- | --- | --- | --- | --- | --- |
| 1 | M | 0.211 | 7.3 | M,N,P | e |
| 4 | L | 0.361 | 6.3 | M,S,L,I,H,F | b |

**CS Score normalized** (1-4= variable, 5= average, 6-9= conserved).
**B/E:** Burried (b) or Exposed (e) residue.

**Table (5):** The *PRSS1* gene functions and its appearance in network and genome.

| Function | FDR | Genes<br>in<br>network | Genes<br>in<br>genome |
| --- | --- | --- | --- |
| serine-type endopeptidase inhibitor activity | 0.001356361 | 4 | 44 |
| renal system process involved in regulation of systemic arterial blood pressure | 0.001356361 | 3 | 13 |
| regulation of hemostasis | 0.001356361 | 4 | 54 |
| regulation of coagulation | 0.001356361 | 4 | 56 |
| regulation of blood coagulation | 0.001356361 | 4 | 54 |
| regulation of wound healing | 0.003496886 | 4 | 74 |
| platelet activation | 0.006856052 | 5 | 211 |
| endopeptidase inhibitor activity | 0.007599469 | 4 | 99 |
| endopeptidase regulator activity | 0.007599469 | 4 | 102 |
| peptidase inhibitor activity | 0.007599469 | 4 | 101 |
| blood microparticle | 0.008676071 | 4 | 108 |
| regulation of endopeptidase activity | 0.009340491 | 5 | 251 |
| regulation of peptidase activity | 0.009856636 | 5 | 258 |
| peptidase regulator activity | 0.013382385 | 4 | 128 |
| renal system process | 0.013666888 | 3 | 45 |
| endocrine process | 0.013666888 | 3 | 42 |
| negative regulation of endopeptidase activity | 0.013666888 | 4 | 140 |
| regulation of blood vessel size | 0.013666888 | 3 | 44 |
| regulation of tube size | 0.013666888 | 3 | 44 |
| negative regulation of peptidase activity | 0.013666888 | 4 | 142 |
| regulation of systemic arterial blood pressure | 0.013666888 | 3 | 43 |
| platelet alpha granule lumen | 0.01586832 | 3 | 48 |
| vascular process in circulatory system | 0.028313203 | 3 | 59 |
| platelet alpha granule | 0.029997483 | 3 | 61 |
| secretory granule lumen | 0.030241005 | 3 | 62 |
| serine-type endopeptidase activity | 0.033116705 | 3 | 67 |
| killing of cells of other organism | 0.033116705 | 2 | 10 |
| exocytosis | 0.033116705 | 4 | 191 |
| disruption of cells of other organism | 0.033116705 | 2 | 10 |
| glomerular filtration | 0.037840129 | 2 | 11 |
| renal filtration | 0.037840129 | 2 | 11 |
| vesicle lumen | 0.040192639 | 3 | 76 |
| enzyme inhibitor activity | 0.040192639 | 4 | 214 |
| protein kinase C-activating G-protein coupled receptor signaling pathway | 0.040192639 | 2 | 12 |
| cytoplasmic membrane-bounded vesicle lumen | 0.040192639 | 3 | 76 |
| positive regulation of collagen metabolic process | 0.044298773 | 2 | 13 |
| positive regulation of collagen biosynthetic process | 0.044298773 | 2 | 13 |
| regulation of blood pressure | 0.044298773 | 3 | 81 |
| platelet degranulation | 0.044771122 | 3 | 82 |
| positive regulation of ERK1 and ERK2 cascade | 0.04525757 | 3 | 83 |
| negative regulation of multicellular organismal process | 0.045315882 | 4 | 230 |
| positive regulation of coagulation | 0.048108433 | 2 | 15 |
| negative regulation of hydrolase activity | 0.048108433 | 4 | 242 |
| collagen metabolic process | 0.048108433 | 3 | 87 |
| positive regulation of hemostasis | 0.048108433 | 2 | 15 |
| positive regulation of blood coagulation | 0.048108433 | 2 | 15 |
| positive regulation of multicellular organismal metabolic process | 0.048108433 | 2 | 15 |
| multicellular organismal macromolecule metabolic process | 0.049591958 | 3 | 91 |
| regulation of endocrine process | 0.051022071 | 2 | 16 |
| regulation of collagen biosynthetic process | 0.051022071 | 2 | 16 |
| tissue homeostasis | 0.051393559 | 3 | 94 |
| zymogen activation | 0.052534202 | 3 | 97 |
| regulation of collagen metabolic process | 0.052534202 | 2 | 17 |
| multicellular organismal metabolic process | 0.052534202 | 3 | 97 |
| positive regulation of release of sequestered calcium ion into cytosol | 0.052534202 | 2 | 17 |
| serine-type peptidase activity | 0.052961891 | 3 | 98 |
| serine hydrolase activity | 0.055058507 | 3 | 101 |
| cellular metal ion homeostasis | 0.055058507 | 4 | 264 |
| positive regulation of Ras protein signal transduction | 0.055058507 | 2 | 18 |
| collagen biosynthetic process | 0.057743853 | 2 | 19 |
| multicellular organismal homeostasis | 0.057743853 | 3 | 105 |
| regulation of multicellular organismal metabolic process | 0.057743853 | 2 | 19 |
| positive regulation of small GTPase mediated signal transduction | 0.057743853 | 2 | 19 |
| cobalamin metabolic process | 0.0619817 | 2 | 20 |
| fibrinolysis | 0.0619817 | 2 | 20 |
| cellular cation homeostasis | 0.063339568 | 4 | 284 |
| regulation of ERK1 and ERK2 cascade | 0.063981924 | 3 | 111 |
| metal ion homeostasis | 0.064852029 | 4 | 288 |
| cellular ion homeostasis | 0.066330871 | 4 | 292 |
| extracellular matrix organization | 0.066330871 | 4 | 292 |
| extracellular structure organization | 0.066330871 | 4 | 293 |
| ERK1 and ERK2 cascade | 0.07491573 | 3 | 120 |
| positive regulation of calcium ion transport into cytosol | 0.094036985 | 2 | 26 |
| positive regulation of cytosolic calcium ion concentration | 0.094313684 | 3 | 131 |

**\*FDR:** false discovery rate is greater than or equal to the probability that this is a false positive.

**Table (6):**
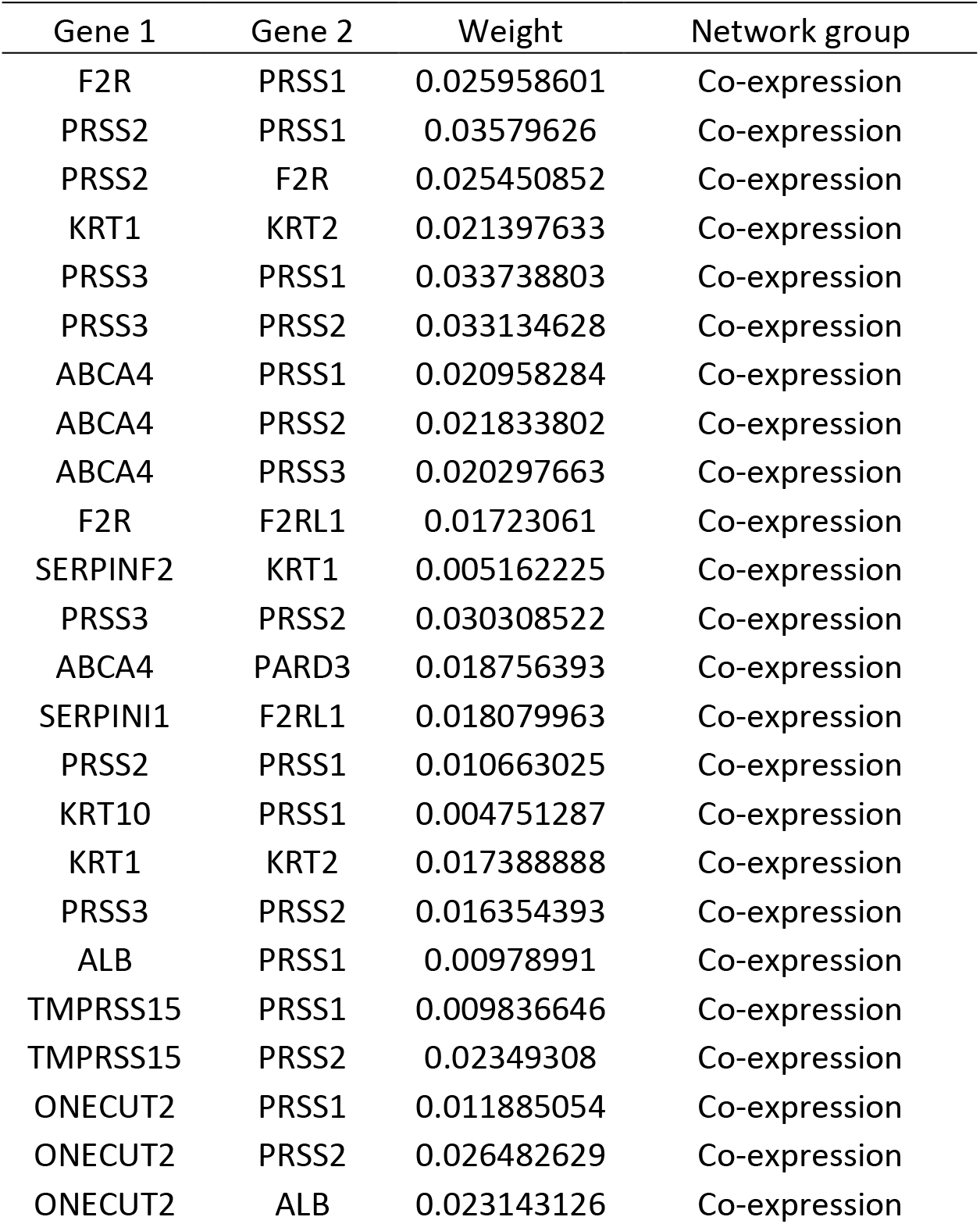
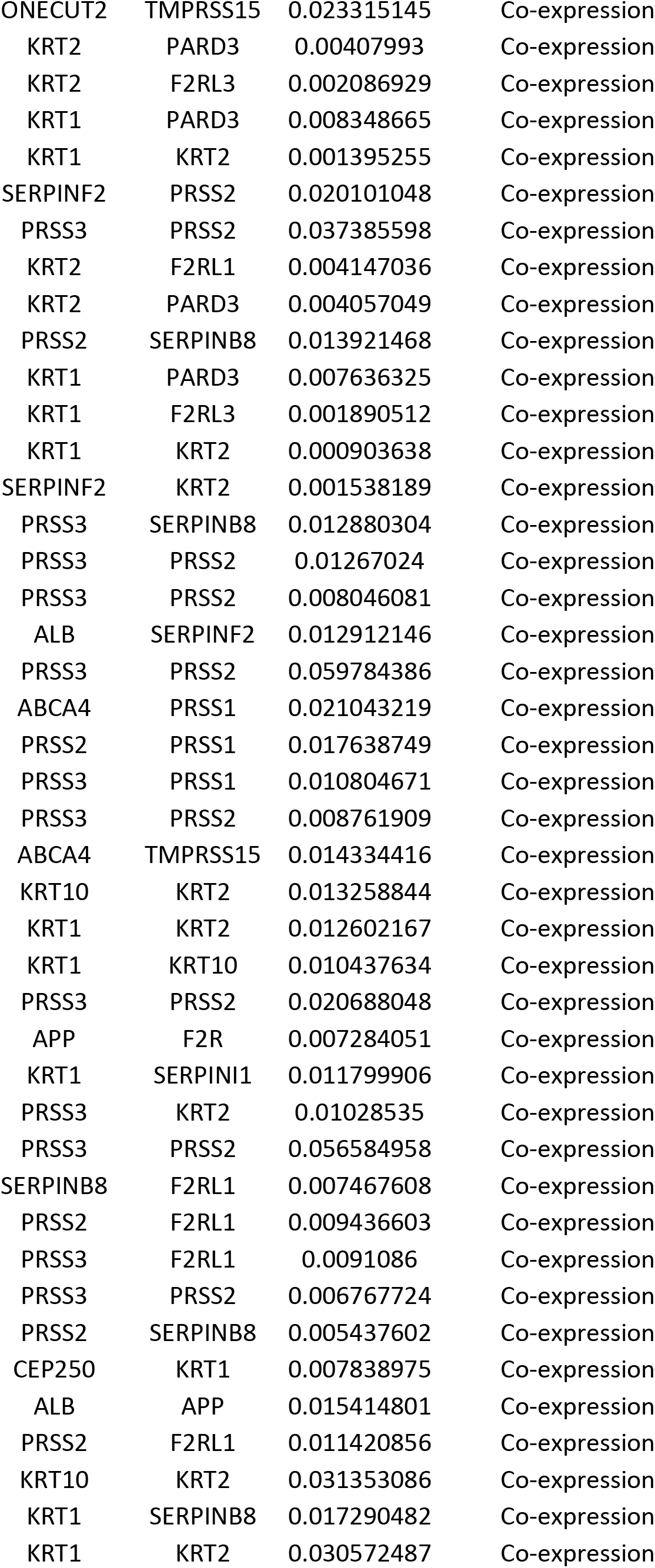
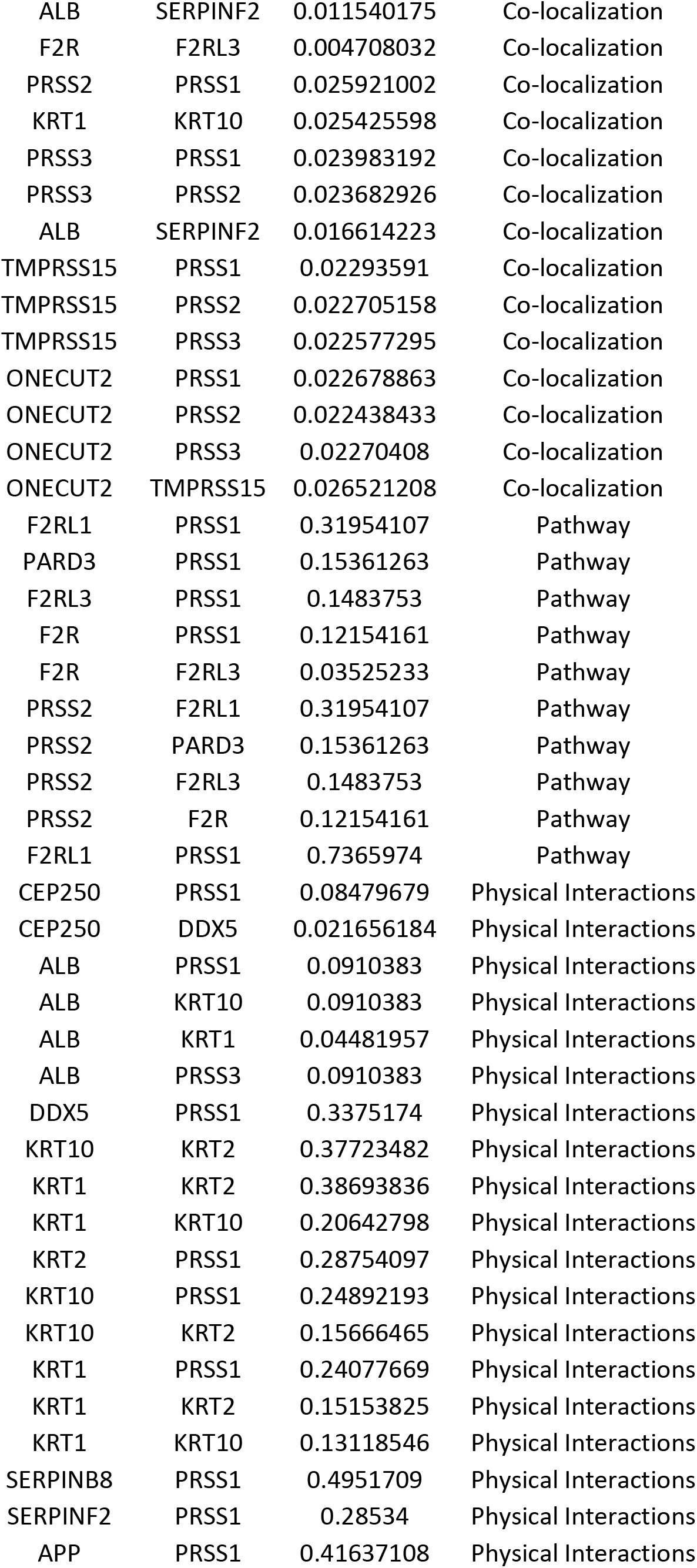
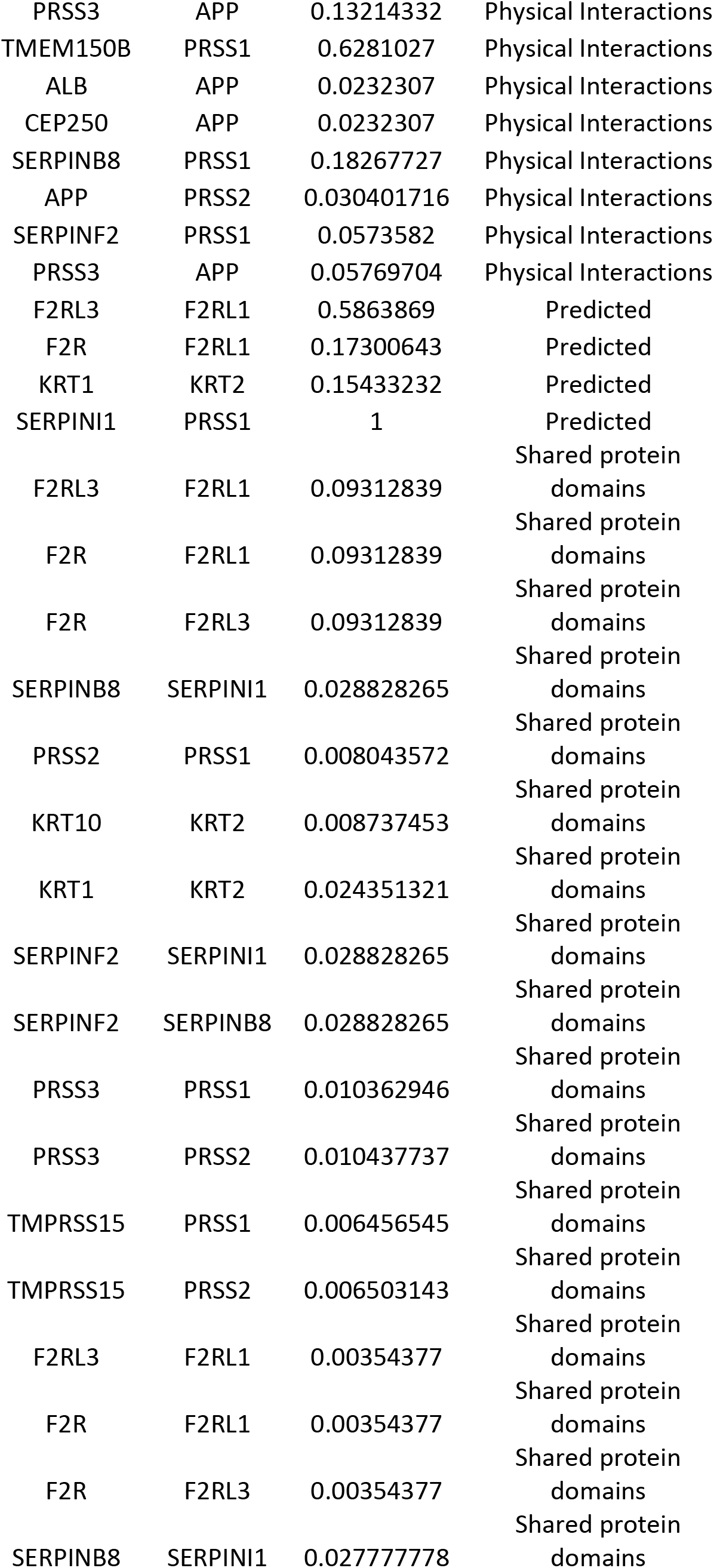
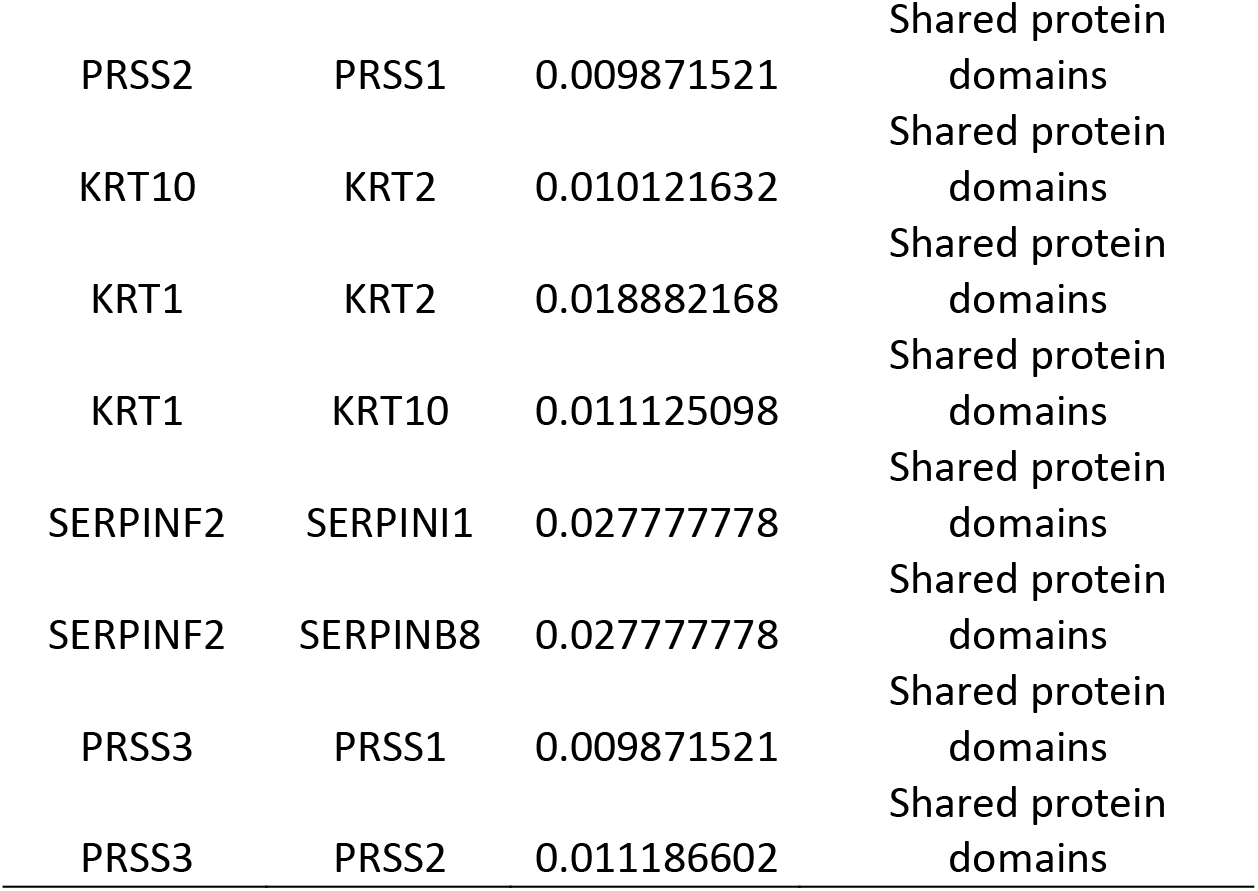
the gene co-expressed, share domain and Interaction with *MEFV* gene network.

In the light of our study we recommend to perform whole exome sequencing on suspicious patients(3, 16, 17) and their parents for pathogenic variant (M1R & L4P) besides the ones that has been screened in other studies (6, 64) to allow early diagnosis and prevention and/or management of symptoms.

## 8. Conclusion

A total of two nsSNPs (M1R & L4P) were predicted to be responsible for the structural and functional modifications of *PRSS1* protein. It is evident from the deep comprehensive in silico analysis that, these two nsSNPs might serve as, a novel biomarkers for the prognosis of HP. These findings may also facilitate the development of novel therapeutic elements for associated diseases.

## Data Availability

All relevant data are within the paper and its supporting information files.

## Acknowledgment

The authors wish to acknowledgment the enthusiastic cooperation of Africa City of Technology - Sudan.

## References

1. Charnley RM. Hereditary pancreatitis. World journal of gastroenterology. 2003;9(1):1–4.

2. Jackson WD. Pancreatitis: etiology, diagnosis, and management. Current opinion in pediatrics. 2001;13(5):447–51.

3. Dai LN, Chen YW, Yan WH, Lu LN, Tao YJ, Cai W. Hereditary pancreatitis of 3 Chinese children: Case report and literature review. Medicine. 2016;95(36):e4604.

4. Le Bodic L, Bignon JD, Raguenes O, Mercier B, Georgelin T, Schnee M, et al. The hereditary pancreatitis gene maps to long arm of chromosome 7. Human molecular genetics. 1996;5(4):549–54.

5. Dyrla P, Nowak T, Gil J, Adamiec C, Bobula M, Saracyn M. [Hereditary pancreatitis]. Polski merkuriusz lekarski: organ Polskiego Towarzystwa Lekarskiego. 2016;40(236):113–6.

6. Rebours V, Levy P, Ruszniewski P. An overview of hereditary pancreatitis. Digestive and liver disease: official journal of the Italian Society of Gastroenterology and the Italian Association for the Study of the Liver. 2012;44(1):8–15.

7. Howes N, Lerch MM, Greenhalf W, Stocken DD, Ellis I, Simon P, et al. Clinical and genetic characteristics of hereditary pancreatitis in Europe. Clinical gastroenterology and hepatology: the official clinical practice journal of the American Gastroenterological Association. 2004;2(3):252–61.

8. Lee SK. [Hereditary pancreatitis]. The Korean journal of gastroenterology = Taehan Sohwagi Hakhoe chi. 2005;46(5):358–67.

9. Raphael KL, Willingham FF. Hereditary pancreatitis: current perspectives. Clinical and experimental gastroenterology. 2016;9:197–207.

10. Masamune A, Kikuta K, Hamada S, Nakano E, Kume K, Inui A, et al. Nationwide survey of hereditary pancreatitis in Japan. Journal of gastroenterology. 2018;53(1):152–60.

11. Mora J, Comas L, Ripoll E, Goncalves P, Queralto JM, Gonzalez-Sastre F, et al. Genetic mutations in a Spanish population with chronic pancreatitis. Pancreatology: official journal of the International Association of Pancreatology (IAP) [et al]. 2009;9(5):644–51.

12. Raty S, Piironen A, Babu M, Pelli H, Sand J, Uotila S, et al. Screening for human cationic trypsinogen (PRSS1) and trypsinogen inhibitor gene (SPINK1) mutations in a Finnish family with hereditary pancreatitis. Scandinavian journal of gastroenterology. 2007;42(8):1000–5.

13. Sibert JR. Hereditary pancreatitis in England and Wales. Journal of medical genetics. 1978;15(3):189–201.

14. Goebell H, Ammann R, Creutzfeldt W. History of the European pancreatic club: the first 40 years 1965-2005. The development of the European pancreatic club as a scientific society. Pancreatology: official journal of the International Association of Pancreatology (IAP) [et al]. 2005;5 Suppl 1:1–15.

15. Applebaum-Shapiro SE, Finch R, Pfutzer RH, Hepp LA, Gates L, Amann S, et al. Hereditary pancreatitis in North America: the Pittsburgh-Midwest Multi-Center Pancreatic Study Group Study. Pancreatology: official journal of the International Association of Pancreatology (IAP) [et al]. 2001;1(5):439–43.

16. Joergensen MT, Brusgaard K, Cruger DG, Gerdes AM, Schaffalitzky de Muckadell OB. Genetic, epidemiological, and clinical aspects of hereditary pancreatitis: a population-based cohort study in Denmark. The American journal of gastroenterology. 2010;105(8):1876–83.

17. Atlas AB, Orenstein SR, Orenstein DM. Pancreatitis in young children with cystic fibrosis. The Journal of pediatrics. 1992;120(5):756–9.

18. Masamune A. Genetics of pancreatitis: the 2014 update. The Tohoku journal of experimental medicine. 2014;232(2):69–77.

19. Langner C. [Hereditary gastric and pancreatic cancer]. Der Pathologe. 2017;38(3):164–9.

20. Lowenfels AB, Maisonneuve P, DiMagno EP, Elitsur Y, Gates LK, Jr., Perrault J, et al. Hereditary pancreatitis and the risk of pancreatic cancer. International Hereditary Pancreatitis Study Group. Journal of the National Cancer Institute. 1997;89(6):442–6.

21. Malka D, Hammel P, Maire F, Rufat P, Madeira I, Pessione F, et al. Risk of pancreatic adenocarcinoma in chronic pancreatitis. Gut. 2002;51(6):849–52.

22. Weiss FU. Pancreatic cancer risk in hereditary pancreatitis. Frontiers in physiology. 2014;5:70.

23. Lowenfels AB, Maisonneuve P, Whitcomb DC, Lerch MM, DiMagno EP. Cigarette smoking as a risk factor for pancreatic cancer in patients with hereditary pancreatitis. Jama. 2001;286(2):169–70.

24. Lowenfels AB, Maisonneuve P, Whitcomb DC. Risk factors for cancer in hereditary pancreatitis. International Hereditary Pancreatitis Study Group. The Medical clinics of North America. 2000;84(3):565–73.

25. Mastoraki A, Tzortzopoulou A, Tsela S, Danias N, Sakorafas G, Smyrniotis V, et al. Hereditary pancreatitis: dilemmas in differential diagnosis and therapeutic approach. Journal of gastrointestinal cancer. 2014;45(1):22–6.

26. Patel MR, Eppolito AL, Willingham FF. Hereditary pancreatitis for the endoscopist. Therapeutic advances in gastroenterology. 2013;6(2):169–79.

27. Kargl S, Kienbauer M, Duba HC, Schofl R, Pumberger W. Therapeutic step-up strategy for management of hereditary pancreatitis in children. Journal of pediatric surgery. 2015;50(4):511–4.

28. LaRusch J, Solomon S, Whitcomb DC. Pancreatitis Overview. In: Adam MP, Ardinger HH, Pagon RA, Wallace SE, Bean LJH, Stephens K, et al., editors. GeneReviews((R)). Seattle (WA): University of Washington, Seattle University of Washington, Seattle. GeneReviews is a registered trademark of the University of Washington, Seattle. All rights reserved.; 1993.

29. Rivera Rivera ED. [Pancreatitis, genes and islet cells auto transplant; updates and new horizons]. Revista de gastroenterologia del Peru: organo oficial de la Sociedad de Gastroenterologia del Peru. 2017;37(2):156–61.

30. Solomon S, Whitcomb DC, LaRusch J. PRSS1-Related Hereditary Pancreatitis. In: Adam MP, Ardinger HH, Pagon RA, Wallace SE, Bean LJH, Stephens K, et al., editors. GeneReviews((R)). Seattle (WA): University of Washington, Seattle University of Washington, Seattle. GeneReviews is a registered trademark of the University of Washington, Seattle. All rights reserved.; 1993.

31. LaRusch J, Barmada MM, Solomon S, Whitcomb DC. Whole exome sequencing identifies multiple, complex etiologies in an idiopathic hereditary pancreatitis kindred. JOP: Journal of the pancreas. 2012;13(3):258–62.

32. Kim JY, Choi SH, Ihm JS, Kim SJ, Kim IJ, Kim CM. [A case of R122H mutation of cationic trypsinogen gene in a pediatric patient with hereditary pancreatitis complicated by pseudocyst and hemosuccus pancreaticus]. The Korean journal of gastroenterology = Taehan Sohwagi Hakhoe chi. 2005;45(2):130–6.

33. Jamer T, Iwanczak B. [Genetic mutations as a cause of acute recurrent pancreatitis in children - case report and literature review]. Developmental period medicine. 2016;20(3):228–34.

34. Lal A, Lal DR. Hereditary pancreatitis. Pediatric surgery international. 2010;26(12):1193–9.

35. Le Marechal C, Masson E, Chen JM, Morel F, Ruszniewski P, Levy P, et al. Hereditary pancreatitis caused by triplication of the trypsinogen locus. Nature genetics. 2006;38(12):1372–4.

36. Simon P, Weiss FU, Sahin-Toth M, Parry M, Nayler O, Lenfers B, et al. Hereditary pancreatitis caused by a novel PRSS1 mutation (Arg-122 --< Cys) that alters autoactivation and autodegradation of cationic trypsinogen. The Journal of biological chemistry. 2002;277(7):5404–10.

37. Nemeth BC, Sahin-Toth M. Human cationic trypsinogen (PRSS1) variants and chronic pancreatitis. American journal of physiology Gastrointestinal and liver physiology. 2014;306(6):G466–73.

38. Chen JM, Le Marechal C, Lucas D, Raguenes O, Ferec C. “Loss of function” mutations in the cationic trypsinogen gene (PRSS1) may act as a protective factor against pancreatitis. Molecular genetics and metabolism. 2003;79(1):67–70.

39. Whitcomb DC, Gorry MC, Preston RA, Furey W, Sossenheimer MJ, Ulrich CD, et al. Hereditary pancreatitis is caused by a mutation in the cationic trypsinogen gene. Nature genetics. 1996;14(2):141–5.

40. Lee TY, Oh HC, Kim MH, Kwon S, Lee SS, Seo DW, et al. [Three cases of hereditary pancreatitis in two households in the same family associated with R122H mutation in cationic trypsinogen gene]. The Korean journal of gastroenterology = Taehan Sohwagi Hakhoe chi. 2007;49(6):395–9.

41. Hanck C, Schneider A, Whitcomb DC. Genetic polymorphisms in alcoholic pancreatitis. Best practice & research Clinical gastroenterology. 2003;17(4):613–23.

42. Chen X, Chang JT. Planning bioinformatics workflows using an expert system. Bioinformatics (Oxford, England). 2017;33(8):1210–5.

43. Bajard A, Chabaud S, Cornu C, Castellan AC, Malik S, Kurbatova P, et al. An in silico approach helped to identify the best experimental design, population, and outcome for future randomized clinical trials. Journal of clinical epidemiology. 2016;69:125–36.

44. Mustafa MI, Abdelhameed T, Abdelrhman F, Osman S, Hassan M. Novel deleterious nsSNPs within MEFV gene that could be used as Diagnostic Markers to Predict Hereditary Familial Mediterranean Fever: Using bioinformatics analysis. bioRxiv. 2018:424796.

45. Benson DA, Karsch-Mizrachi I, Clark K, Lipman DJ, Ostell J, Sayers EW. GenBank. Nucleic acids research. 2012;40(Database issue):D48–53.

46. Artimo P, Jonnalagedda M, Arnold K, Baratin D, Csardi G, de Castro E, et al. ExPASy: SIB bioinformatics resource portal. Nucleic acids research. 2012;40(Web Server issue):W597–603.

47. Sim NL, Kumar P, Hu J, Henikoff S, Schneider G, Ng PC. SIFT web server: predicting effects of amino acid substitutions on proteins. Nucleic acids research. 2012;40(Web Server issue):W452–7.

48. Capriotti E, Altman RB. Improving the prediction of disease-related variants using protein three¬dimensional structure. BMC bioinformatics. 2011;12 Suppl 4:S3.

49. Choi Y, Sims GE, Murphy S, Miller JR, Chan AP. Predicting the functional effect of amino acid substitutions and indels. PloS one. 2012;7(10):e46688.

50. Hecht M, Bromberg Y, Rost B. Better prediction of functional effects for sequence variants. BMC genomics. 2015;16 Suppl 8:S1.

51. Calabrese R, Capriotti E, Fariselli P, Martelli PL, Casadio R. Functional annotations improve the predictive score of human disease-related mutations in proteins. Human mutation. 2009;30(8):1237–44.

52. Lopez-Ferrando V, Gazzo A, de la Cruz X, Orozco M, Gelpi JL. PMut: a web-based tool for the annotation of pathological variants on proteins, 2017 update. Nucleic acids research. 2017;45(W1):W222–w8.

53. Capriotti E, Fariselli P, Casadio R. I-Mutant2.0: predicting stability changes upon mutation from the protein sequence or structure. Nucleic acids research. 2005;33(Web Server issue):W306–10.

54. Cheng J, Randall A, Baldi P. Prediction of protein stability changes for single-site mutations using support vector machines. Proteins. 2006;62(4):1125–32.

55. Ashkenazy H, Erez E, Martz E, Pupko T, Ben-Tal N. ConSurf 2010: calculating evolutionary conservation in sequence and structure of proteins and nucleic acids. Nucleic acids research. 2010;38(Web Server issue):W529–33.

56. Ashkenazy H, Abadi S, Martz E, Chay O, Mayrose I, Pupko T, et al. ConSurf 2016: an improved methodology to estimate and visualize evolutionary conservation in macromolecules. Nucleic acids research. 2016;44(W1):W344–50.

57. Oni OO, Owoade AA, Adeyefa CAO. Design and evaluation of primer pairs for efficient detection of avian rotavirus. Tropical animal health and production. 2018;50(2):267–73.

58. Warde-Farley D, Donaldson SL, Comes O, Zuberi K, Badrawi R, Chao P, et al. The GeneMANIA prediction server: biological network integration for gene prioritization and predicting gene function. Nucleic acids research. 2010;38(Web Server issue):W214–20.

59. Meyer MJ, Lapcevic R, Romero AE, Yoon M, Das J, Beltran JF, et al. mutation3D: Cancer Gene Prediction Through Atomic Clustering of Coding Variants in the Structural Proteome. Human mutation. 2016;37(5):447–56.

60. Pettersen EF, Goddard TD, Huang CC, Couch GS, Greenblatt DM, Meng EC, et al. UCSF Chimera--a visualization system for exploratory research and analysis. Journal of computational chemistry. 2004;25(13):1605–12.

61. George Priya Doss C, Sudandiradoss C, Rajasekaran R, Choudhury P, Sinha P, Hota P, et al. Applications of computational algorithm tools to identify functional SNPs. Functional & integrative genomics. 2008;8(4):309–16.

62. Chandrasekaran G, Hwang EC, Kang TW, Kwon DD, Park K, Lee JJ, et al. In silico analysis of the deleterious nsSNPs (missense) in the homeobox domain of human HOXB13 gene responsible for hereditary prostate cancer. 2017;90(2):188–99.

63. Kosaloglu Z, Bitzer J, Halama N, Huang Z, Zapatka M, Schneeweiss A, et al. In silico SNP analysis of the breast cancer antigen NY-BR-1. BMC cancer. 2016;16(1):901.

64. Oracz G, Kolodziejczyk E, Sobczynska-Tomaszewska A, Wejnarska K, Dadalski M, Grabarczyk AM, et al. The clinical course of hereditary pancreatitis in children - A comprehensive analysis of 41 cases. Pancreatology: official journal of the International Association of Pancreatology (IAP) [et al]. 2016;16(4):535–41.

